# Exploration of Sensory and Spinal Neurons Expressing GRP in Itch and Pain

**DOI:** 10.1101/472886

**Authors:** Devin M. Barry, Xue-Ting Liu, Qianyi Yang, Xian-Yu Liu, Xiansi Zeng, Ben-Long Liu, Zhou-Feng Chen

**Affiliations:** Center for the Study of Itch, Washington University School of Medicine, St. Louis, MO 63110, U.S.A.; Department of Anesthesiology, Washington University School of Medicine, St. Louis, MO 63110, U.S.A.; Department of Psychiatry, Washington University School of Medicine, St. Louis, MO 63110, U.S.A.; Developmental Biology, Washington University School of Medicine, St. Louis, MO 63110, U.S.A.; The Second Affiliated Hospital, The State Key Laboratory of Respiratory Disease, Guangdong Provincial Key Laboratory of Allergy & Clinical Immunology, Sino-French Hoffmann Institute, Center for Immunology, Inflammation, & Immune-mediated disease, Guangzhou Medical University, Guangzhou, Guangdong 510260, P.R. China; College of Life Sciences, Xinyang Normal University, 237 Nanhu Road, Xinyang 464000, P. R. China.

## Abstract

Gastrin-releasing peptide (GRP) is a putative itch-specific neurotransmitter, but definite evidence in the dorsal root ganglion (DRG) and spinal cord is lacking. We generated and validated a *Grp*-Cre knock-in (*Grp*^Cre-KI^) mouse line whereby *Grp* neurons are genetically labeled. Cre-dependent marking analysis revealed exclusive innervation of the upper epidermis of the skin by GRP fibers. Importantly, optical stimulation of *Grp* fibers expressing channel rhodopsin (ChR2) in the skin evoked itch but not pain-related scratching behaviors, while conditional deletion of *Grp* in sensory neurons attenuated non-histaminergic itch. In contrast, intersectional genetic ablation of spinal *Grp* neurons did not affect itch nor pain transmission. Our study demonstrates a role of GRP in sensory neurons in itch and suggests that GRP sensory neurons are dedicated to itch transmission. *Grp*^Cre-KI^ mice provide a long-sought avenue for investigating peripheral coding mechanism of itch and further interrogation of itch-nerve fibers in the skin under chronic pruritus.

**Highlights:**

- Validated expression of a *Grp*-Cre knock-in line in sensory neurons that innervate the skin
- Opto-activation of *Grp* sensory neurons evokes itch behavior
- Conditional deletion of *Grp* in sensory neurons reduces non-histaminergic itch behavior
- Intersectional ablation of *Grp* spinal neurons does not affect itch or pain behaviors

## Introduction

Primary afferents of sensory neurons release fast neurotransmitters (e.g. glutamate) and neuropeptides to activate postsynaptic receptors in the spinal cord dorsal horn to transmit itch and pain information (Barry et al., 2017; Bautista et al., 2014; Braz et al., 2014; Jeffry et al., 2011; McNeil and Dong, 2012). A fundamental challenge for elucidating the coding mechanisms of itch vs. pain is to identify itch-specific neural circuits. Since 2009, a number of studies have used genetic and/or chemical inactivation approaches to identify itch-specific neurons (Bourane et al., 2015; Han et al., 2013; Mishra and Hoon, 2013; Sun et al., 2009). One limitation of these approaches, however, is that the role of ablated neurons in itch/pain might have been masked due to compensation. This is especially a concern for DRG neurons, where presence of nociceptors is believed to be overwhelming, which is a basis for the population coding theory (LaMotte et al., 2014; Ma, 2012). One way to circumvent the problem and/or complement the loss-of-function approach is to activate the neurons of interest followed by characterizing the nature of evoked behaviors (e.g. scratching behavior) which may reflect either itch, pain or both (Shimada and LaMotte, 2008).

Gastrin releasing peptide (GRP) has been suggested to be a principal neuropeptide in transmitting itch from sensory neuron primary afferents to GRP-receptor (GRPR) neurons in the superficial laminae of the dorsal horn (Sun and Chen, 2007). The potential evolutionary root for GRP as a generic itch transmitter has recently been further supported by the finding that GRP in the suprachiasmatic nucleus of hypothalamus is crucial for contagious itch behavior (Yu et al., 2017) as well as its conserved expression in the nervous system of rodents and primates (Nattkemper et al., 2013; Takanami et al., 2016; Takanami et al., 2014). Moreover, studies suggest that GRP primarily transmits non-histaminergic itch as well as long-lasting itch information to the spinal cord (Akiyama et al., 2013; Barry et al., 2017; Nattkemper et al., 2013; Zhao et al., 2013; Zhao et al., 2014). However, there are considerable disagreements on the expression of *Grp* mRNA in DRGs (Barry et al., 2016; Mishra and Hoon, 2013; Solorzano et al., 2015), which could be ascribed to the inherent problem of BAC-based transgenic *Grp*-eGFP mouse lines(Gong et al., 2003) as well as technical caveats (Barry et al., 2016). Regardless of whether GRP is expressed in DRGs, the direct experimental evidence for the role of GRP in DRGs is lacking(Braz et al., 2014). On the other hand, there are conflicting results concerning the role of *Grp* spinal neurons in itch and pain. One study argued that *Grp* spinal neurons are a relaying station for itch-specific transmission (Mishra and Hoon, 2013), whereas the other claimed that they constitute a “leaky gate” for itch and pain (Sun et al., 2017). While we failed to find significant level of GRP protein which can be attributable to endogenous expression from *Grp* spinal neurons (Barry et al., 2016; Zhao et al., 2013), the possibility that GRP peptide released and rapidly degraded from *Grp* spinal neurons cannot be excluded for certainty.

Because BAC-transgenic *Grp*-eGFP mice fail to express eGFP in DRGs (Mishra and Hoon, 2013; Solorzano et al., 2015; Sun et al., 2017), it is imperative to tag the *Grp* locus with a genetic marker using a gene targeting approach, thereby enabling GRP sensory neurons accessible for functional studies. To this aim, we generated and validated a long-sought knock-in mouse line in which Cre recombinase (Cre) is driven under the control of the endogenous *Grp* promoter (*Grp*^Cre-KI^) and a floxed *Grp* allele for conditional deletion. In addition, we used an intersectional genetic approach to investigate the role of *Grp* spinal neurons in itch and pain.

## Results

### *Grp*^Cre-KI^ Expression in Sensory Neurons and Skin Fibers

To investigate the role of *Grp* sensory neurons in itch, a targeting construct was generated with an eGFP-Cre recombinase cassette and knocked-in at exon 1 start codon of *Grp* (*Grp*^Cre-KI^) in mice (**Figure 1A; Figures S1A and S1B**). eGFP-positive neurons in DRG were detected following immunohistochemistry (IHC) with a GFP antibody, but epifluorescent eGFP signals were not observed (**Figures 1B and 1C**). Moreover, the GFP antibody did not detect any signals in *Grp*^WT^ tissues (**Figure S1C**). To validate specific expression of *Grp* in DRG neurons, we employed RNAScope *in situ* hybridization (ISH) methods (Wang et al., 2012) and found that *Grp* is detected in a subset of wild type DRG neurons (~7%) but not in *Grp* KO tissues (**Figures 1D and 1E**). This result is consistent with *Grp* mRNA (~7%) detected by conventional digoxygenin (DIG) (**Figure 1F**) as well as we previously reported (Barry et al., 2016; Zhao et al., 2013). To further confirm specific expression of *Cre* in DRG neurons of *Grp*^Cre-KI^ mice, double ISH was performed with *Grp* and *Cre* probes. We found that nearly all *Cre*^+^ neurons co-expressed *Grp* (258 of 275 *Cre* neurons) and, likewise, almost all *Grp*^+^ neurons co-expressed *Cre* (258 of 278 *Grp* neurons) (**Figures 1G and 1H**). Moreover, *Cre* was not observed in *Grp*^WT^ mice (**Figures S1D and S1E**).

**Figure 1.**
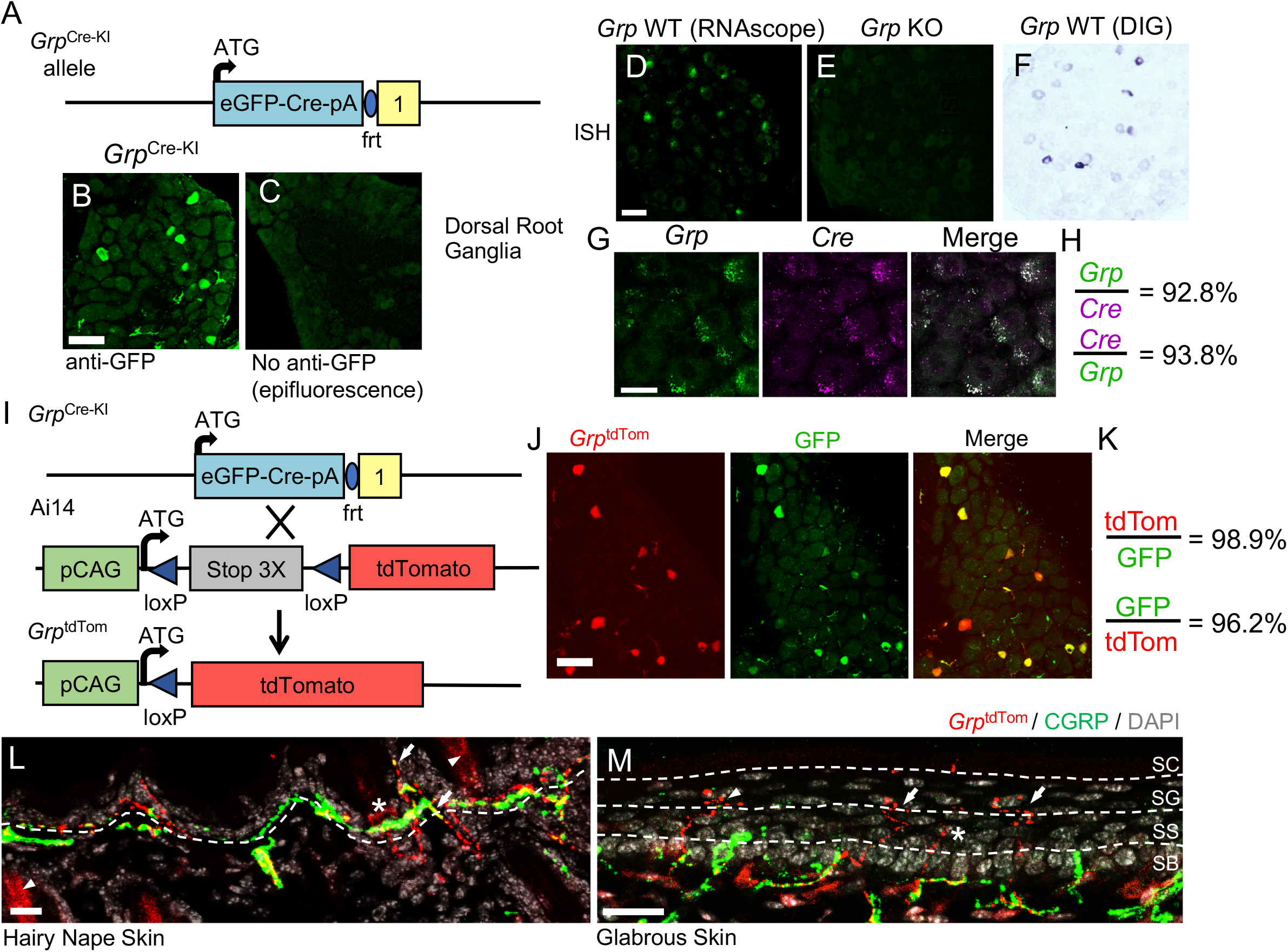
Validated expression of *Grp* and *Cre* in *Grp*^Cre-KI^ Sensory Neurons. (**A**) Schematic of eGFP-Cre Neo cassette in *Grp* allele to generate *Grp*^Cre-KI^ mice. (**B**) eGFP IHC from *Grp*^Cre-KI^ DRG section. (**C**) Image of eGFP epifluorescent signal (no GFP antibody) in *Grp*^Cre-KI^ DRG. Scale bar in **B**, 50 μm. (**D**) RNAscope method of *Grp* ISH in WT DRG. Scale bar in **D**, 50 μm. (**E**) *Grp* ISH in *Grp* KO DRG. (**F**) *Grp* IsH using conventional DIG method in WT DRG. (**G**) ISH images of *Grp* and *Cre* in *Grp*^Cre-KI^ DRG. Scale bar in **G**, 20 μm. (**H**) Percentage of *Cre*- or Grp-expressing neurons that express *Grp* or *Cre*, respectively. (**I**) Schematic of *Grp*^Cre-KI^ mating with Ai9 reporter line to produce *Grp*^tdTom^ mice. (**J**) IHC images of tdTomato and eGFP in *Grp*^tdTom^ DRG. Scale bar in **J**, 50 μm. (**K**) Percentage of eGFP^+^ neurons co-expressing tdTomato or of tdTomato^+^ neurons co-expressing eGFP. (**L**) IHC image of tdTomato, CGRP and DAPI in *Grp*^tdTom^ nape skin. Arrows indicate apparent *Grp*^tdTom^ fibers coexpressing CGRP. Arrowheads indicate autofluorescent hair shafts. Dashed line indicates epidermal/dermal boundary. (**M**) IHC images of tdTomato, CGRP, and DAPI in glabrous paw skin with stratum basilis (SB), stratum spinosum (SS), stratum granulosum (SG), and stratum corneum (SC) epidermal layers marked by dashed lines. Arrows indicate *Grp*^tdTom^ fibers with ‘S’ or ‘Z’ pattern nerve endings. Arrowhead indicates *Grp*^tdTom^ fibers with apparent bush endings. Scale bar in **L**, **M**, 20 μm. n = 3 mice and 10 sections for **G**, **H**, **J** and **K**.

To characterize innervation of *Grp* sensory neurons in the skin, *Grp*^Cre-KI^ mice were crossed with the tdTomato flox-stop reporter line (Ai14) (Madisen et al., 2010) to generate mice with tdTomato expression in *Grp*^+^ neurons (*Grp*^tdTom^)(**Figure 1I**). To validate tdTomato expression, we performed IHC of eGFP with tdTomato epifluorescent signals in *Grp*^tdTom^ DRGs. tdTomato was expressed in nearly all DRG neurons that expressed eGFP in *Grp*^tdTom^ mice (98/99 tdTom/eGFP and 98/102 eGFP/tdTom neurons) (**Figure 1J and 1K**). tdTomato expression was not detected in DRG of *Grp*^WT^;Ai14 mice (**Figure S2A**). The results indicate that tdTomato expression faithfully recapitulates *Grp* expression in DRG neurons and can be used as a surrogate for *Grp* fibers. To characterize the innervation organization of GRP fibers in the skin, we examined a wide array of tissues and found that they received innervation from *Grp*^tdTom^ fibers with a focus on the epidermis and cutaneous structures (**Table S1**). *Grp*^tdTom^ was observed in nerve fibers of hairy nape skin, with apparent co-expression in some fibers expressing calcitonin gene-related peptide (CGRP), a peptidergic marker of nociceptive neurons (**Figure 1L, arrows**) (Zylka et al., 2005). Some *Grp*^tdTom^ fibers that were CGRP-negative wrapped around the upper epidermal regions of hair follicles as apparent circular or penetrating follicle neck endings (**Figure 1L, asterisk**). *Grp*^tdTom^ fibers showed apparent intertwining with CGRP fibers or with some fibers showing co-expression. Expression of tdTomato was not observed in the skin of *Grp*^WT^;Ai14 mice (**Figure S2B**). Next, we examined the thicker glabrous skin of the paw to better visualize the termination zones of *Grp*^tdTom^ fibers. Many *Grp*^tdTom^ fibers projected from the dermal/epidermal boundary mostly straight up through the stratum basilis and stratum spinosum, but then meandered and wrapped around keratinocytes in apparent ‘S’ or ‘Z’ pattern free endings (arrows) or as bush/cluster endings (arrowhead) within the stratum granulosum and terminated ~5 μm from the stratum corneum (**Figure 1M**, red). In contrast, most of the CGRP fibers along with a few *Grp*^tdTom^ fibers (asterisk) were straight with less complexity and terminated mostly in the stratum spinosum (**Figure 1M**, green). *Grp*^tdTom^ was not co-expressed with myelinated fibers using an antibody against neurofilament heavy (NF-H)(**Figures S2C-S2E**) (Lawson and Waddell, 1991; Price, 1985), and no *Grp*^tdTom^ fibers were observed in any sensory structures that are typically innervated by myelinated fibers including Merkel cell-complexes, Meissner corpuscles, vibrissa follicle-sinus complexes, or sebaceous glands (**Table S1**). *Grp*^tdTom^ fibers were also absent from, heart, esophagus, intestine, kidney, liver, lung and testis (**Table S1**). *Grp*^tdTom^ fibers were rarely observed in the epithelial layers of the tongue, cornea and bladder, (**Figures S2F-S2H**). *Grp*^tdTom^ fibers were also barely present in skeletal muscle within coursing nerve bundles with CGRP fibers but did not innervate muscle fibers or the neuromuscular junction (**Figure S2I and S2J**).

### *Grp* Is Expressed in a Subset of Peptidergic and Non-Peptidergic Sensory Neurons

DRG neurons can be extensively divided into different subsets based upon expression profiles of various molecular markers as well as their size distributions(Basbaum et al., 2009; Snider and McMahon, 1998). We analyzed *Grp*^tdTom^ expression in DRG using several classic neuronal markers. *Grp*^tdTom^ sensory neurons co-expressed CGRP (arrowheads) (69%, 76/113) and/or the non-peptidergic Isolectin B4 (IB4)-binding (arrowheads)(55.8%, 61/113)(**Figures 2A and 2B**) (Hunt and Rossi, 1985; Silverman and Kruger, 1990). Moreover, many *Grp*^tdTom^ neurons coexpressed both CGRP and IB4 (arrows)(36.3%, 41/113)(**Figures 2A and 2B**). Most *Grp*^tdTom^-CGRP^+^ neurons expressed relatively low levels of CGRP based upon staining intensities with few expressing high levels (McCoy et al., 2013; McCoy et al., 2012; Schutz et al., 2004). *Grp*^tdTom^ neurons also co-expressed transient receptor potential cation channel subfamily V member 1 (TRPV1) (Caterina et al., 1997; Shim et al., 2007) (arrowheads) (74.8%, 86/115)(**Figure 2C and 2D**). Consistent with our finding that *Grp*^tdTom^ was not observed in myelinated fibers of skin sensory structures, *Grp*^tdTom^ DRG neurons rarely expressed NF-H (2.4%, 5/204) (**Figures S3A and S3B**). Next we performed ISH to determine expression of *Grp* with an itch-specific receptor *Mrgpra3* (Liu et al., 2009), or with histamine receptor 1, *Hrh1*(Han et al., 2006). *Mrgpra3* was co-expressed in most *Grp* neurons (65.2%, 47/72)(**Figures 2E and 2F**), and nearly half of *Mrgpra3* neurons coexpressed *Grp* (41.2%, 47/114). *Hrh1* was co-expressed in a majority of *Grp* neurons (56.2% 36/64 neurons)(**Figures 2G and 2H**), whereas a small portion of *Hrh1* neurons co-expressed *Grp* (20.8%, 36/173). Lastly, we further characterized *Grp*^tdTom^ DRG neurons by measuring the area (μm^2^) of perikarya as well as for CGRP and IB4 neurons (**Figure S3C**). The areas of *Grp*^tdTom^ neuron perikarya were typically less than that of CGRP and IB4 neurons (*Grp*^tdTom^ 292 ± 13 μm^2^ vs CGRP 369 ± 16 μm^2^ vs IB4 364 ± 9 μm^2^). Overall the size profiles indicate that *Grp*^tdTom^ neurons are mostly smaller sized neurons and show similar distributions to CGRP or IB4 populations, albeit shifted to a slightly smaller area profile than CGRP or IB4.

**Figure 2.**
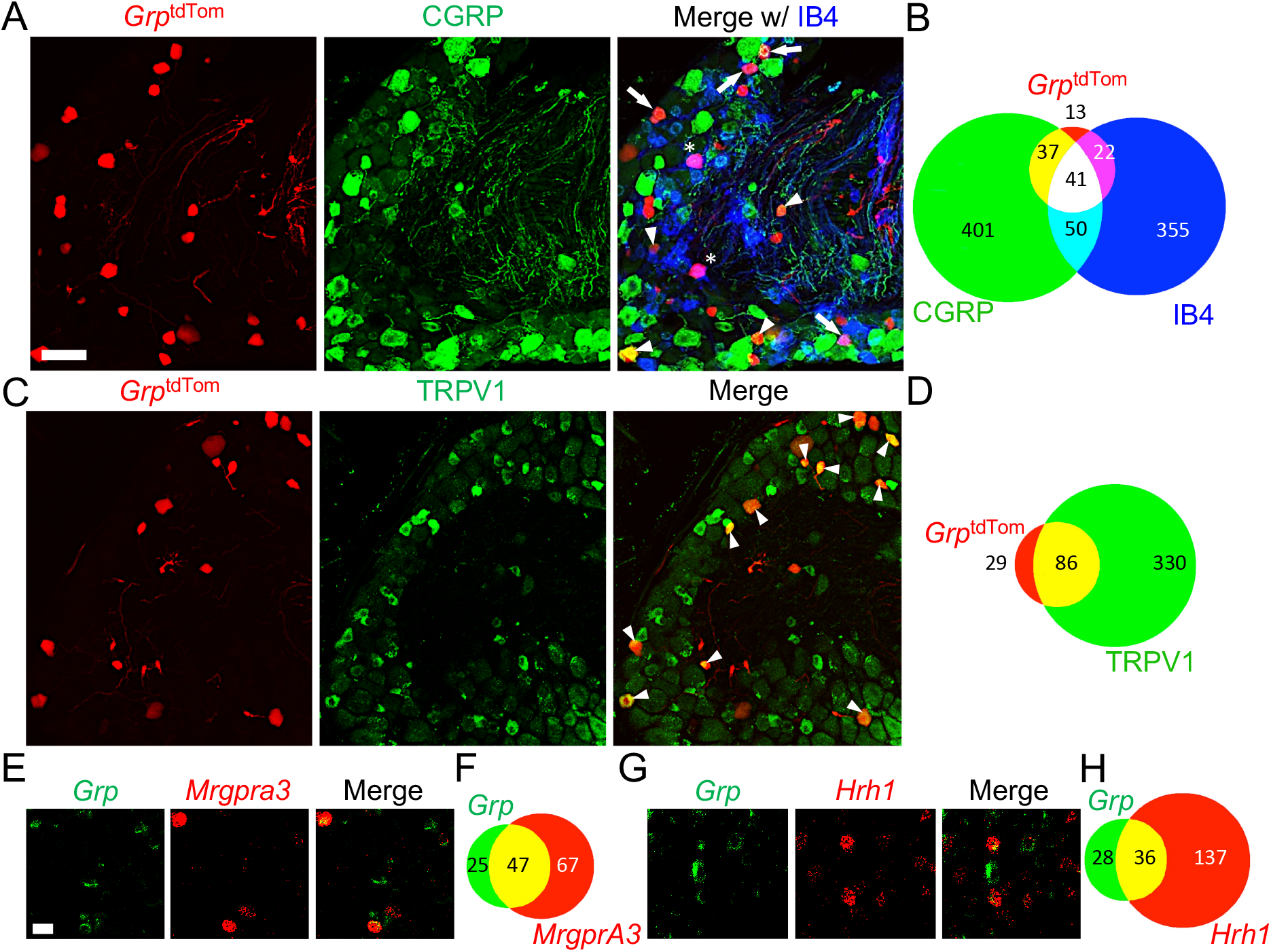
*Grp* expression is enriched in a subset of sensory neurons with both peptidergic and non-peptidergic markers. (**A**) IHC image of tdTomato, CGRP and w IB4 in *Grp*^tdTom^ DRG. Arrows indicate tdTomato neurons co-expressing CGRP and IB4-binding. Arrowheads indicate tdTomato neurons expressing CGRP only. Asterisks indicate tdTomato neurons with IB4-binding only. Scale bar in **A**, 50 μm. (**B**) Venn Diagram of DRG neurons with tdTomato, CGRP expression and IB4-binding from *Grp*^tdTom^ mice. (**C**) IHC of tdTomato and TRPV1 in *Grp*^tdTom^ DRG. Arrowheads indicate tdTomato neurons expressing TRPV1. (**D**) Diagram of DRG neurons with tdTomato and TRVP1 expression. (**E**) ISH of *Grp* and *Mrgpra3* in WT DRG. Scale bar in **E**, 20 μm. (**F**) Diagram of DRG neurons with *Grp* and *Mrgpra3* expression. (**G**) ISH of *Grp* and *Hrh1* in WT DRG. (**H**) Diagram of DRG neurons with *Grp* and *Hrh1* expression, n = 3 mice and 9 sections.

### *Grp* Sensory Neurons Are Activated by Pruritogens

To investigate the responsiveness of *Grp* sensory neurons to pruritogens, we performed Ca^2+^ imaging of cultured DRG neurons from *Grp*^tdTom^ mice. tdTomato^+^ neurons were observed in culture and loaded with the ratiometric indicator fura2-AM for live-cell imaging of intracellular Ca^2+^ levels as described (Kim et al., 2016)(**Figures 3A and 3B**). Chloroquine (CQ) and histamine (Hist), two archetypal pruritogens for nonhistaminergic and histaminergic itch, respectively, were applied separately, followed by capsaicin (Cap), a potent agonist of TRPV1 (Caterina et al., 2000) which can also signal itch via Mrpgra3 neurons(Han et al., 2012), and potassium chloride (KCl) as a positive control response (**Figures 3C and 3D**). We found that ~74% of *Grp*^tdTom^ neurons responded to CQ (72/97) and ~52% of CQ-responsive neurons expressed *Grp*^tdTom^ (72/138) (**Figure 3E**). ~52% *Grp*^tdTom^ neurons responded to Hist (51/97) and ~33% of Hist-responsive neurons expressed *Grp*^tdTom^ (51/156)(**Figure 3E**)., ~41% of *Grp*^tdTom^ neurons were responsive to both CQ and Hist (40/97)(**Figure 3E**). Most *Grp*^tdTom^ neurons also responded to Cap (~81%, 79/97) (**Figure 3E**). As with DRG tissues, cultured *Grp*^tdTom^ DRG neurons co-expressed CGRP, IB4 and TRPV1 but almost no co-expression with NF-H (**Figures S4A and S4B**). Taken together, the data suggested that the vast majority of *Grp*^tdTom^ sensory neurons are activated by pruritogens.

**Figure 3.**
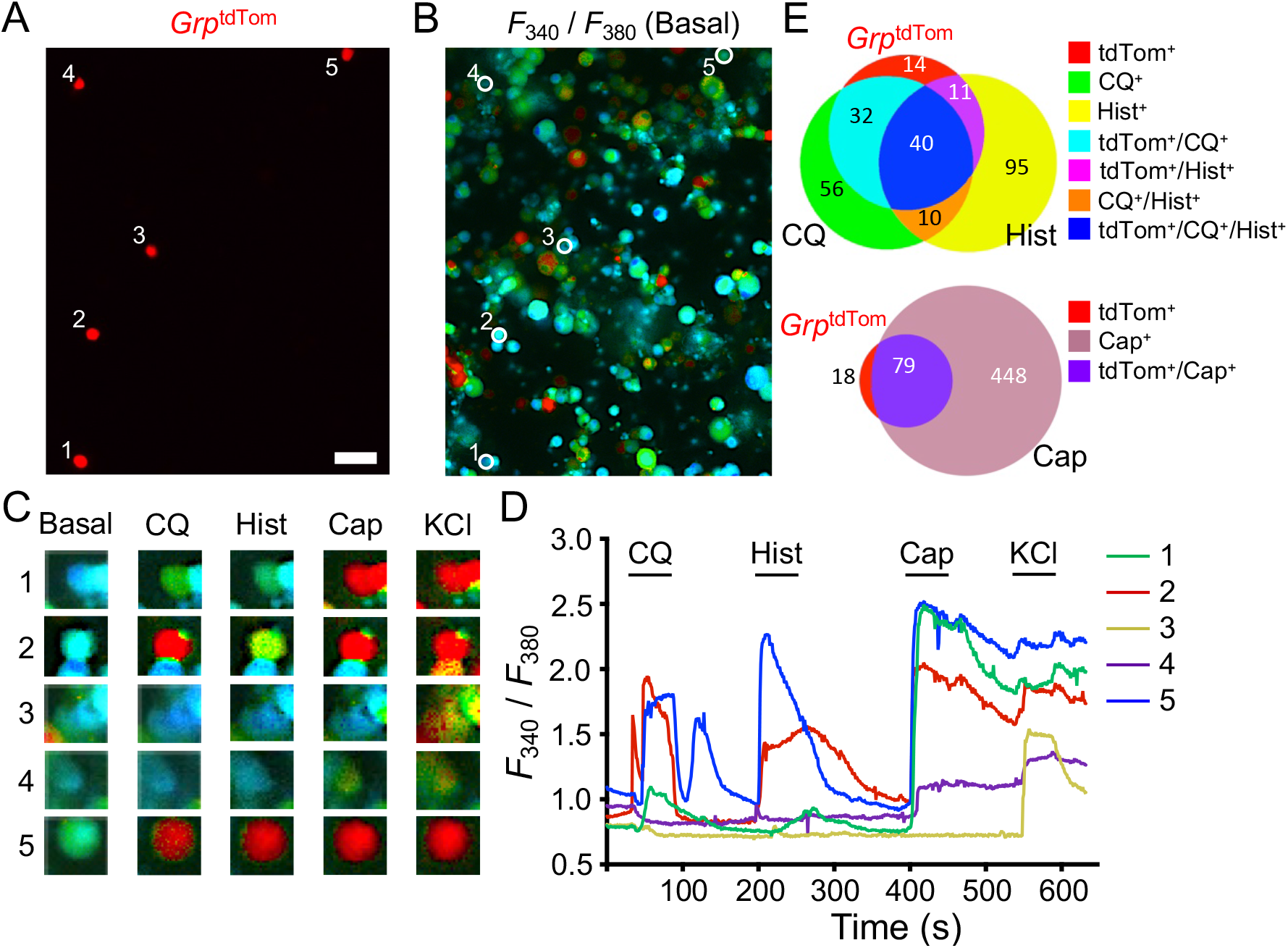
Pruritogens induce Ca^2+^ responses in *Grp*^+^ sensory neurons. (**A, B**) Image of tdTomato neurons (**A**) and *F*_340_ / *F*_380_ signal (**B**) from *Grp*^tdTom^ DRG cultures loaded with fura 2-AM. Scale bar in *A*, 50 μm. (**C**) Snapshots of *Grp*^tdTom^ neuron intracellular Ca^2+^ levels at basal and during CQ (1 mM), Hist (100 μM), Cap (100 nM) or KCI (30 mM) applications. (**D**) *F*_340_ / *F*_380_ traces from *Grp*^tdTom^ neurons with application of CQ, Hist, Cap and KCI. (**E**) Venn diagram of *Grp*^tdTom^ DRG neurons responsive to CQ, Hist or Cap and/or expressing tdTomato. n = 4 mice and 1424 neurons for **A-E**.

### Opto-Activation of *Grp* Sensory Neuron Fibers Induces Itch-Specific Behavior

Since *Grp* is also expressed in the spinal cord, it is technically difficult to inactivate *Grp* neurons exclusively in DRGs. To overcome the problem, we crossed *Grp*^Cre-KI^ mice with a flox-stop channel rhodopsin-eYFP (ChR2-eYFP) line (Ai32) (Madisen et al., 2012) to generate mice with ChR2-eYFP expression in *Grp* neurons (*Grp*^ChR2^)(**Figures 4A and 4B**), so that *Grp* sensory neurons can be opto-activated to assess their role in itch and pain. As with *Grp*^tdTom^, ChR2-eYFP was detected in both CGRP^+^ and IB4-binding sensory neurons in *Grp*^ChR2^ mice (**Figure S5A**), whereas ChR2-eYFP was absent in *Grp*^WT^;Ai32 mice (**Figure S5B**). eYFP was also detected in nerve fibers of the nape skin and cheek skin that expressed βIII-tubulin (**Figure 4C; Figure S5D**), but no ChR2-eYFP was detected in *Grp*^WT^;Ai32 mice (**Figure S5C**). We also detected *Grp*^ChR2^ fibers in the glabrous skin that appeared intertwined or co-expressed with some CGRP^+^ fibers in both the dermal and epidermal layers (**Figure S5E**). Moreover, DiI injection was injected into the nape or cheek skin to retrogradely label DRG neurons (Zele et al., 2010) that confirmed *Grp*^ChR2^ DRG neurons innervate the nape skin (**Figure S5F**), as well as *Grp*^ChR2^ TG neurons that innervated the cheek skin (**Figure S5G**).

**Figure 4.**
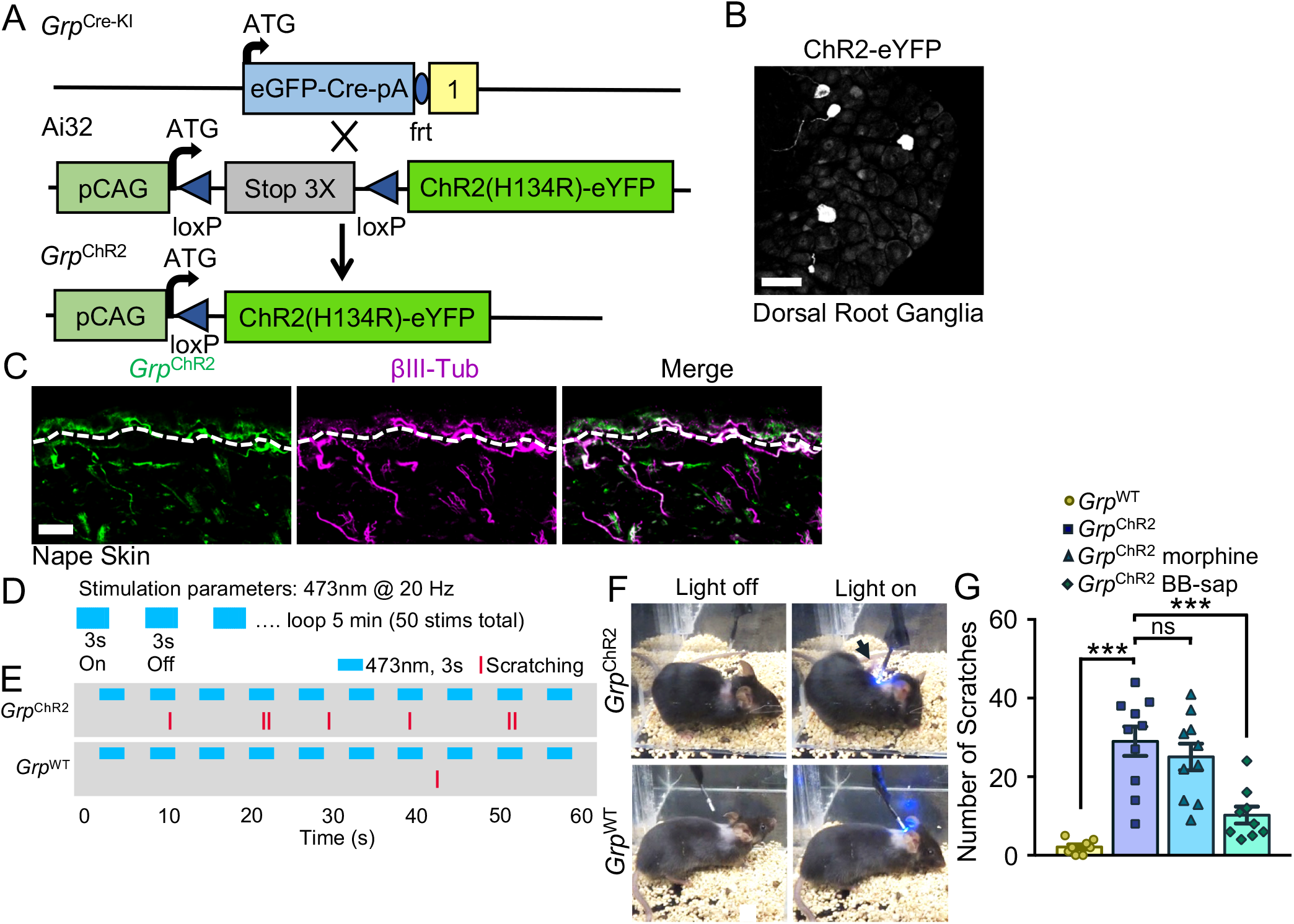
Opto-activation of *Grp*^+^ sensory neuron skin fibers evokes itch behavior. (**A**) Schematic of *Grp*^Cre KI^ mating with Ai32 ChR2-eYFP line to produce *Grp*^ChR2^ mice. (**B**) IHC Image of eYFP expression in *Grp*^ChR2^ DRG. Scale bar in *B*, 50 μm. (**C**) IHC images of eYFP and ßlll-Tubulin in *Grp*^ChR2^ nape skin. Dashed lines indicate epidermal/dermal boundary. Scale bar in *C*, 100 μm. (**D**) Optical parameters of skin fiber stimulation *Grp*^ChR2^ and *Grp*^WT^ mice. (**E**) Raster plot of scratching behavior induced by light stimulation of skin in *Grp*^ChR2^ and *Grp*^WT^ mice. (**F**) Snapshots of *Grp*^ChR2^ and *Grp*^WT^ mice with light off or on. Arrow indicates hind paw scratching the nape when light is on. (**G**) Total number of scratches during 5-min light stimulation experiment in *Grp*^WT^, *Grp*^ChR2^, *Grp*^ChR2^ morphine-treated and *Grp*^ChR2^ BB-sap-treated mice. (**H**) n = 8 – 10 mice, one-way ANOVA with Tukey *post hoc*, *** *p* < 0.001, ns – not significant.

To assess whether activation of *Grp* sensory neurons transmits itch-specific information, we took advantage of *Grp* fiber innervation in the skin and observed corresponding behaviors upon optogenetic activation. *Grp*^ChR2^ mice were stimulated with 473 nm blue light with a fiber optic held just above the nape skin (15 mW power from fiber tip) at 1, 5, 10 or 20 Hz with a 3 s On-Off cycle for 5 min total (**Figure 4D**). Notably, *Grp*^ChR2^ mice exhibited significant scratching behaviors during blue light stimulation trials at 10 and 20 Hz, whereas *Grp*^WT^ mice showed almost no scratching (**Figures 4E and 4F; Figure S6A; Movies S1 and S2**). For the 3 s stimulation trials that induced scratching, the latency to scratch was ~1.7 s (**Figure S6B**). At 20 Hz, the percentage of trials that induced scratching in *Grp*^ChR2^ mice was much greater compared to *Grp*^WT^ mice (43.2% vs 2.5%, *p* < 0.001)(**Figure S6BC**). Similarly, the total number of scratches induced during the 5 min test at 20 Hz was significant compared to *Grp*^WT^ mice (24.3 vs 1.9 scratches, *p* < 0.001)(**Figure 4G**). To determine whether the blue light-induced scratching behaviors were indication of pain or itch, we treated *Grp*^ChR2^ mice with systemic morphine (5 mg/kg intraperitoneal) for analgesia since morphine induced scratching and analgesia are mediated by distinct pathways in the spinal cord(Liu et al., 2011b). We found that morphine analgesia in *Grp*^ChR2^ mice had no significant effect on scratching induced by 20 Hz blue-light stimulation (**Figure 4G**) or on the percentage of trials inducing scratching behavior (**Figure S6C**). This finding suggested that scratching is likely to be itch-related rather than pain. To confirm this, intrathecal injection (i.t.) of bombesin-saporin (BB-sap, 200 ng) was performed to ablate GRPR neurons in the spinal cord and block itch transmission (Sun et al., 2009). Indeed, *Grp*^ChR2^ mice treated with BB-sap showed a significant reduction in scratching (24.3 vs 7.9 scratches, *p* < 0.001)(**Figure 4G**) as well as the percentage of trials inducing scratching behaviors at 20 Hz (43.2% vs 14.4%, *p* < 0.001)(**Figure S6C**). Lastly, we monitored spontaneous scratching behaviors for 30 min prior to and 30 min following the 5 min-20 Hz stimulation test and found a significant increase in scratching following stimulation in *Grp*^ChR2^ mice compared to *Grp*^WT^ (68.4 vs 13.9 scratches, *p* < 0.001), which was not affected by morphine analgesia but was almost completely blocked in BB-sap treated mice (68.4 vs 10.1, *p* < 0.001)(**Figure S6D**). These data indicate that activation of *Grp* sensory neurons selectively transmits itch sensation.

### Conditional Deletion of *Grp* in Sensory Neurons Attenuates Itch Behavior

Previous evidence from global *Grp* KO mice indicated that loss of GRP attenuated non-histaminergic itch behavior, whereas histaminergic itch and pain transmission were normal(Wan et al., 2017). However, it remained unclear whether the source of GRP from sensory neurons, spinal neurons or the brain were important for itch. To determine the role of GRP in sensory neurons, we generated a floxed *Grp* allele and crossed it with the sensory neuron specific Cre-recombinase line, Na_v_1.8^Cre^ (Stirling et al., 2005), to conditionally delete *Grp* expression in sensory neurons (**Figure 5A**). *Grp* expression was mostly absent in DRG of *Grp*^F/F^;Na_v_1.8^Cre^ mice compared to *Grp*^F/F^ control littermates (**Figures 5B and 5C**), but *Grp* expression in dorsal horn was not affected (**Figures S7A and S7B**). Next, we performed acute itch behavior tests and found significant reductions of scratching behavior in *Grp*^F/F^;Na_v_1.8^Cre^ mice compared to *Grp*^F/F^ control for the non-histaminergic pruritogens CQ (240 ± 24 vs 143 ± 16 scratches, *p* < 0.01)(**Figure 5D**), the PAR2 agonist SLIGRL-NH_2_ (Shimada et al., 2006) (222 ± 17 vs 109 ± 23 scratches, *p* < 0.01)(**Figure 5E**), and the MrgprC11 agonist BAM8-22 (Liu et al., 2011a) (46 ± 4 vs 29 ± 3 scratches, *p* < 0.01)(**Figure 5F**). However, scratching behavior for histamine was similar in *Grp*^F/F^;Na_v_1.8^Cre^ mice compared to *Grp*^F/F^ control (53 ± 6 vs 51 ± 7 scratches, *p* = 0.79) (**Figure 5G**). Taken together, these results indicate that GRP in sensory neurons is important for non-histaminergic itch transmission but not for histaminergic itch.

**Figure 5.**
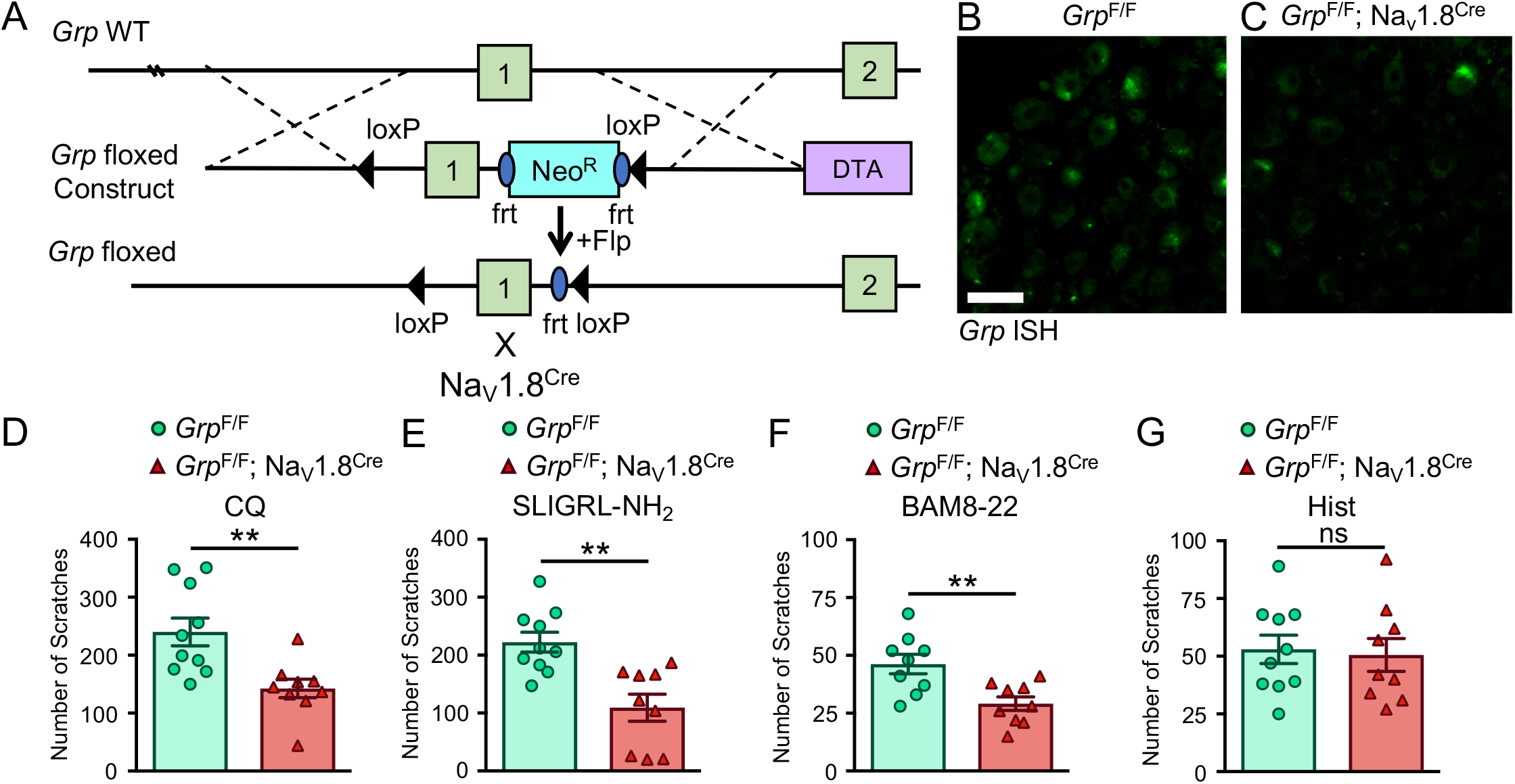
Attenuated itch behaviors in mice with conditional deletion of *Grp* in sensory neurons. (**A**) Schematic of targeting strategy for inserting loxP sites into *Grp* allele to generate *Grp* floxed (*Grp*^F/F^) mice and subsequent crossing with Na_v_1.8^Cre^ mice. (**B, C**) ISH images of *Grp* expression in DRG sections from (**B**) *Grp*^F/F^ and (**C**) *Grp*^F/F^; Na_v_1.8^Cre^ mice. Scale bar in **B**, 50 μm. (**D - G**) Mean number of scratches induced by i.d. nape injection of (**D**) CQ (200 μg), (**E**) SLIGRL-NH_2_ (100 μg), (**F**) BAM8-22 (100 μg) and (**G**) Hist (200 μg) in *Grp*^F/F^ and *Grp*^F/F^; Na_v_1.8^Cre^ littermates. n = 9 – 10 mice, unpaired *t* test in **D - G**, ** *p* < 0.01.

### Intersectional Ablation Reveals that *Grp* Spinal Neurons Are Dispensable for Itch and Pain behaviors

To test the role of *Grp* spinal interneurons in transmitting itch and pain behaviors while not affecting *Grp* expressing neurons in the DRG and brain, we used an intersectional genetic approach (Bourane et al., 2015; Duan et al., 2014) to ablate *Grp* neurons specifically in the dorsal spinal cord and spinal trigeminal nucleus subcaudalis (SpVc). *Grp*^Cre-KI^ mice were crossed with the *Lbx1^Flpo^* and *Tau^ds-DTR^* lines to express the diphtheria toxin receptor (DTR) exclusively in *Grp* spinal neurons and mice were injected with diphtheria toxin (DTX) to specifically ablate *Grp* spinal neurons (**Figure 6A**). We validated the *Grp*^Cre-KI^ expression by *Grp* ISH in *Grp*^tdTom^ spinal neurons and found that 89% of tdTomato^+^ neurons express Grp mRNA, and importantly, 93% of Grp mRNA+ neurons express tdTomato (**Figures S8A-C**) We tested the efficiency of DTR-mediated ablation in the spinal cord by *Grp* ISH (**Figures 6B and 6C**) as well as *Grp*^tdTom^ neurons (**Figures 6D and 6E**). Compared to control littermates, DTX injection resulted in ablation of ~82% of *Grp* neurons and ~93% of *Grp*^tdTom^ dorsal horn neurons in ablated mice (**Figures 6F and 6G**).

**Figure 6.**
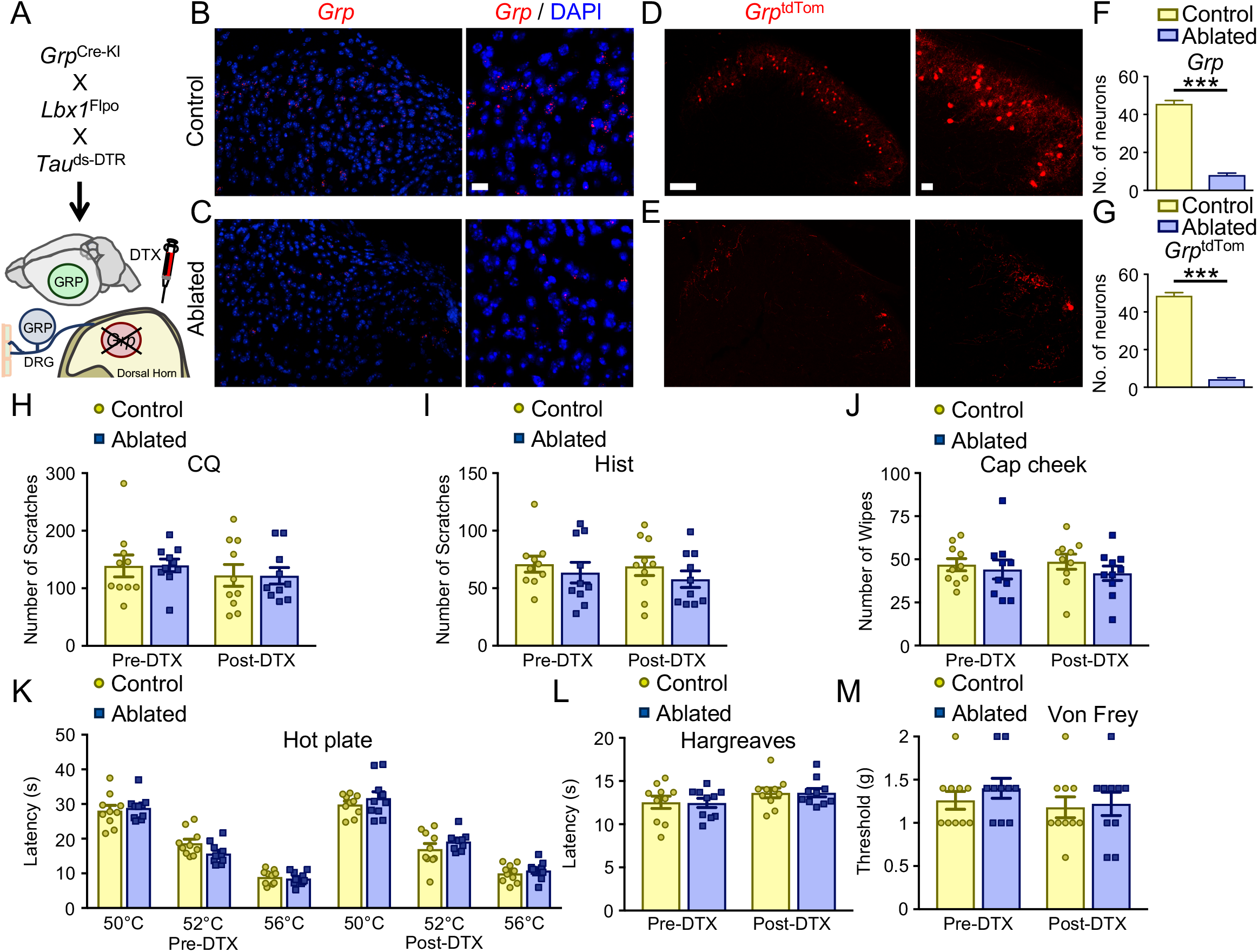
Intersectional ablation of *Grp* spinal neurons does not affect itch or pain behavior responses. (**A**) Schematic of mating strategy of *Grp*^Cre-KI^ with *Lbx1*^Flpo^ and *Tau*^ds-DTR^ lines to generate DTR-expressing *Grp* spinal neurons and subsequent DTX injection to ablate *Grp* spinal neurons. (**B, C**) ISH images of *Grp* expression in cervical spinal cord sections from (**B**) control and (**C**) ablated littermates. Scale bars in **B**, 100 μm and 20 μm. (**D, E**) Epifluorescent images of tdTomato in cervical spinal cord sections from (**D**) *Grp*^tdTom^ control and (**E**) *Grp*^tdTom^ ablated mice. Scale bars in **D**, 100 μm and 20 μm. (**F**) Mean number of *Grp* neurons in control and ablated sections. (**G**) Mean number of *Grp*^tdTom^ neurons in control and ablated sections. n = 3 mice and 10 sections, unpaired *t* test in F and G, *** *p* < 0.001. (**H, I**) Mean number of scratches induced by i.d. nape injection of (**H**) CQ (100 μg) and (**I**) Hist (200 μg) in control and ablated mice pre- and post-DTX injection. (**J**) Mean number of wipes induced by i.d. cheek injection of Cap (20 μg) in control and ablated mice pre- and post-DTX injection. (**K**) Mean response latencies for hot plate assay at 50°C, 52°C and 56°C in control and ablated mice pre- and post-DTX injection. (**L**) Mean response latencies for Hargreaves hind-paw assay in control and ablated mice pre- and post-DTX injection. (**M**) Withdrawal thresholds (in grams of force, g) for Von Frey filament assay in control and ablated mice pre- and post-DTX injection. n = 10 mice, two-way RM ANOVA with Tukey *post-hoc* in **H - M**.

Importantly, we confirmed that *Grp*^tdTom^ neurons in the DRG and brain (cingulate cortex, hippocampus) were not ablated following DTX injection (**Figure S8D**). We analyzed spinal interneuron populations expressing PKC*_γ_*, Pax2, *Grpr* found similar numbers of neurons in control and ablated mice (**Figures S8E and S8F**). *Npr1* neurons were reduced slightly but did not reach statistical significance (77± 3 vs 68 ± 3 neurons, *p* = 0.07) (**Figures S8E and S8F**). This was supported by single cell RT-PCR results showing 25% co-expression of *Npr1* or 10% *Grpr* in *Grp*^tdTom^ neurons (**Table S2**) as well as no co-expression of *Grp*^tdTom^ in PKC*_γ_* cell bodies (**Figure S8E**), and is consistent with recent studies using single nucleus RNA sequencing (snRNA-seq) of spinal cord neurons (Sathyamurthy et al., 2018). To determine the role of spinal *Grp* neurons in itch and pain transmission, we tested acute itch and pain behaviors pre- and post-DTX injection in control and ablated mice. Scratching behaviors for CQ and histamine were similar prior to and following DTX injection in control and ablated mice with no significant differences (**Figures 6H and 6I**). Our finding is in accordance with the observation that few *Grp* neurons seldom expressed c-Fos in response to CQ injection (Bell et al., 2016). Wiping behaviors for capsaicin cheek injection were also comparable between control and ablated mice both pre- and post-DTX injection (**Figure 6J**). Thermal response latencies for hot plate and Hargreaves hind-paw assay were comparable in control and ablated mice both pre- and post-DTX injection, as well as no changes in mechanical thresholds by von Frey filament assay (**Figures 6K-6M**). Taken together, our results indicate that specific ablation of *Grp* spinal neurons does not significantly affect itch or pain behavior responses, suggesting that *Grp* spinal neurons are dispensable for itch and pain transmission.

## Discussion

In this study, we have generated and validated *Grp* and *Cre* expression in sensory neurons of *Grp*^Cre-KI^ line and confirmed that *Cre* recapitulates endogenous *Grp* expression in DRGs. Notably, the finding that the percentage of *Grp* neurons marked by a genetic marker is in accordance with results previously reported using either ISH or IHC approaches (Barry et al., 2016; Panula et al., 1983; Sun and Chen, 2007; Takanami et al., 2014; Zhao et al., 2013). These results confirm the lack of fidelity of a widely used BAC transgenic *Grp* mouse line in sensory neurons. Importantly, the present study demonstrates the role of GRP in sensory neurons for CQ but not histamine-evoked itch behaviors, which are in agreement with previous studies using global *Grp* KO mice(Sun and Chen, 2007; Sun et al., 2009), whereas *Grp* or *Grp* neurons in the spinal cord has no role in CQ or histamine itch. The matching phenotype between GRP global and conditional KO mice, along with dose-dependent scratching behavior induced by i.t. GRP, provide strong evidence for GRP to act as an itch-specific neuropeptide from primary afferents.

Our examination of genetically tagged *Grp* nerve endings reveals extensive innervation organization of pruriceptor fibers highly enriched in the epidermis. Notably, *Grp* is enriched in a unique subset of DRG neurons with both peptidergic and non-peptidergic features that are similar to *Mrgpra3* neurons (Han et al., 2013). Despite co-expression of CGRP in the cell bodies of some *Grp* neurons, *Grp* epidermal fibers mostly lack or do not express detectable levels of CGRP, indicating differences in expression/trafficking of CGRP between cell body and free nerve endings. Moreover, the morphological characteristics of *Grp* epidermal fibers and their elaborate innervation patterns in the stratum granulosum are consistent with a non-peptidergic profile that was previously described in *Mrgprd* fibers (Zylka et al., 2005). The observation that a small percentage of *Grp* neurons gave rise to extensive branching in the skin is in shape contrast to a progressively restriction of CGRP expression in the terminals. The broad innervation of *Grp* fibers may enable their detection of a wide array of pruritogenic stimuli in the skin. The present findings also complement previous studies showing that *Mrgpra3* neurons are dedicated to itch transmission using cell ablation approach (Han et al., 2013). Recent findings that *Nppb* and *Mrgpra3* are not co-expressed in sensory neurons (Huang et al., 2018), as supported by our studies (see also the companion paper on BNP signaling) failed to explain how *Mrgpra3* fibers transmit CQ-evoked itch information. Our observation that more than 60% of *Mrgpra3* express GRP are in support of previous studies indicating that *Mrgpra3* afferents form synaptic contacts with GRPR neurons and use GRP to transmit CQ as well as other nonhistaminergic itch (Han et al., 2013; Liu et al., 2009; Sun et al., 2009). In line with this, both GRP and *Mrgpra3* are up-regulated in DRGs of mice with chronic itch (Zhao et al., 2013). It is worth noting that not all *Grp* neurons were responsive to CQ, histamine and capsaicin, as measured by calcium imaging studies. These non-responding neurons may express receptors for pruritogens which were not investigated in our study. Given increasing numbers of receptors/TRP channels in DRGs have been implicated in itch (Feng et al., 2017; Liu et al., 2012; Oetjen et al., 2017; Steinhoff et al., 2003; Wilson et al., 2013), it is likely that modality specificity is in part encoded through the differential activation and release of myriad neuropeptides. Whether a broad innervation of the epidermis by a small subset of pruriceptors may reflect a signature of modality-specific sensory neurons remains to be determined.

The observation that optogenetic activation of the peripheral nerve endings of *Grp* sensory neurons in the skin nape induced itch-related scratching behaviors provides strong evidence indicating that *Grp* neurons are itch-specific, and functionally match their central target neurons, GRPR neurons, in the spinal cord(Sun et al., 2009; Zhao et al., 2013). A cheek model has been proposed as an excellent approach to differentiate itch vs. pain behavior (Shimada and LaMotte, 2008). However, it is challenging to perform optogenetic stimulation of the nerve endings in the cheek skin of free moving animals as they tend to avoid blue light stimulation when directed at their face. The use the light to stimulate the nape skin is advantageous because under natural environment the application avoids direct contacts between blue light and eyes. Our approach also found that higher frequency (≥ 10Hz) stimulation can more efficiently evoke robust scratching behaviors, whereas previous studies only tested 1 or 5 Hz, which resulted in minimal scratching over the course of 5 minutes (Sun et al., 2017). To the best of our knowledge, the present study is the first demonstration of combinatorial use of optogenetic activation, pain inhibition and neuronal ablation in identification of a subset of itch-specific neurons in DRGs. Our strategy as described here can be applied to differentiate between itch- and pain-related scratching behavior evoked by the blue light onto the nape skin and thus to identify itch- or pain-related primary afferents in sensory neurons.

Our results from conditional deletion of *Grp* specifically in sensory neurons confirm previous reports from global KO mice that indicate GRP largely mediates non-histaminergic itch from the periphery to the spinal cord, whereas NMB primarily transmits histaminergic itch (Wan et al., 2017; Zhao et al., 2013). These results are also in line with the previous results that GRPR mediates non-histaminergic itch while NMBR mediates histaminergic itch (Sun and Chen, 2007; Zhao et al., 2014). The present finding suggests that *Grp* spinal neurons either fail to express GRP protein or GRP protein endogenously expressed in the spinal cord is dispensable for itch (Barry et al., 2016).

As aforementioned, much of the recent controversy on *Grp* expression in sensory neurons and spinal cord could be attributed to low fidelity of Bac-based *Grp*-eGFP or *Grp*-Cre lines generated by GENSAT. Although approximately 90 – 93% of eGFP or Cre neurons express *Grp* mRNA, these lines capture only about 64 – 68% of endogenous *Grp* mRNA cells (Solorzano et al., 2015; Sun et al., 2017). In contrast, we found that approximately 93% of *Grp* spinal neurons express tdTomato, demonstrating a high fidelity of *Grp*^Cre-KI^ mice. A high fidelity of a Cre line is particularly important for interpretation of ablation results. Sun *et al*. reported the impaired itch and pain phenotype of global ablation of *Grp* neurons using BAC-transgenic *Grp*-Cre line (Sun et al., 2017). However, several confounding factors could explain the abnormal phenotype observed. First, the global ablation of *Grp*-Cre neurons in the brain would ablate many *Grp*-Cre neurons in the ACC or mPFC and thus compromise itch/pain transmission. Second, we and the others found that *Grp* neurons normally do not express PKC*_γ_* (Gutierrez-Mecinas et al., 2014). However, despite observing ~10% overlap in expression of *Grp*-Cre with PKC*_γ_*, *Sun et al*. ablated ~ 50% of PKC*_γ_* neurons required for mechanical pain (Malmberg et al., 1997), indicating extensive ectopic ablation of the dorsal horn neurons (Sun et al., 2017). By contrast, our intersectional strategy did not impact PKC*_γ_* expression. Given the contrasting low vs. high level of *Grp* mRNA transcripts in DRG vs. spinal cord, our data are consistent with the notion that there is no correlation between the mRNA/protein level and phenotype at a steady-state or normal physiological condition (Liu et al., 2016). Another caveat is that a neuron could express myriad peptides/receptors. Conversely, a peptide/receptor could be expressed across multiple modality-specific circuits. For example, kappa or mu opioid receptors are expressed in both subpopulations of GRPR neurons as well as in nociceptive neurons in the spinal cord (Liu et al., 2011b; Munanairi et al., 2018). Indeed, our single cell PCR analysis indicates that *Grp* spinal neurons are rather heterogeneous, comprising both itch- and pain-related genes, including NMBR(Zhao et al., 2014), GRPR and somatostatin (Duan et al., 2014). Ablation of a small subset of pruriceptive or nociceptive neurons may not affect overall itch/pain transmission owning to compensatory effect or insufficient ablation. Nevertheless, *Grp*^Cre-KI^ mice could serve as an invaluable genetic resource for investigating the peripheral mechanisms underlying the coding of itch as well as the phenotypic change consequent to the development of chronic itch.

## Experimental Procedures

### Mice

Mice between 7 and 12 weeks old were used for experiments. Mice were housed in clear plastic cages with no more than 5 mice per cage in a controlled environment at a constant temperature of ~23°C and humidity of 50 ± 10% with food and water available *ad libitum*. The animal room was on a 12/12 h light / dark cycle with lights on at 7 am. All experiments were performed in accordance with the guidelines of the National Institutes of Health and the International Association for the Study of Pain and were approved by the Animal Studies Committee at Washington University School of Medicine.

### *In situ* hybridization (ISH)

Conventional ISH was performed as previously described (Barry et al., 2016) using a digoxigenin-labeled cRNA (Roche) antisense probe for *Grp* (bases 149-707 of *Grp* mRNA, NCBI accession NM_175012.4). RNAScope ISH (Advanced Cell Diagnostics, Inc.) was performed as previously described (Wang et al., 2012) using the RNAScope Multiplex Fluorescent Reagent Kit v2 User Manual for Fixed Frozen Tissue. Following probes were used: *Grp* (Mm-Grp-C1, *GenBank:* NM_175012.4, target region bases 22 – 825, Cat No. 317861), *Cre* (Cre-O1-C3, *GenBank:* NC_005856.1, target region bases 2 – 972, Cat No. 474001-C3), *Grpr* (Mm-Grpr-C1, *GenBank:* NM_008177.2, target region bases 463-1596, Cat No. 317871) or *Npr1* (Mm-Npr1-C1, *GenBank:* NM_008727.5, target region bases 941-1882, Cat No. 484531).

### Immunohistochemistry

IHC was performed as previously described (Barry et al., 2016). The following primary antibodies were used: rabbit anti-GFP (1:1000, Molecular Probes, A11122), chicken anti-GFP (1:500, Aves Labs, GFP-1020), rabbit anti-CGRP (1:3000, MilliporeSigma, AB15360), guinea-pig anti-TRPV1 (1:800, Neuromics, GP14100), chicken anti-NF-H (1:2000, EnCor Biotechnology, CPCA-NF-H), rabbit anti-βIII-Tubulin (1:2000, Biolegend, 802001), rabbit anti-PKC*_γ_* (1:1000, Santa Cruz Biotechnology, SC-211), rabbit anti-Pax2 (1:300, ThermoFisher Scientific, 71-6000) IB4-binding was performed using an IB4-AlexaFluor 647 conjugate (2 μg / mL, ThermoFisher Scientific, I32450) and DAPI (2 μg / mL, Sigma, D9542) was used as a nuclear stain of cells.

### DRG culture and Ca2+ imaging

Ca2+ imaging of DRG cultured neurons from *Grp*^tdTom^ mice was performed as previously described (Kim et al., 2016).

### Acute Itch Behavior

Behavioral experiments were performed during the day (0800 – 1500 h) as previously described(Wan et al., 2017). The videos were played back on a computer and quantified by an observer who was blinded to the treatment or genotype. A scratch was defined as a lifting of the hind limb towards the nape or head to scratch and then a replacing of the limb back to the floor, regardless of how many scratching strokes take place between lifting and lowering of the hind limb. Only scratches to the injection site were counted for 30 min.

### Diphtheria toxin receptor-mediated ablation of *Grp* spinal neurons

To ablate DTR-expressing *Grp* spinal neurons for behavioral and histochemical studies, mice were intraperitoneally injected with diphtheria toxin (DTX, 40μg/kg; Sigma Cat. No. D0564) dissolved in saline at day 1 and then again at day 4. Behavioral, ISH and IHC experiments were performed 2 weeks after DTX injection.

### Statistics

Statistical methods are indicated when used. Values are reported as the mean ± standard error of the mean (SEM). Statistical analyses were performed using Prism 7 (v7.0d, GraphPad, San Diego, CA). Normality and equal variance tests were performed for all statistical analyses. *P* < 0.05 was considered statistically significant.

## Acknowledgments

We thank P. Yeager for technical support and lab members for comments. We also thank M. Goulding for generously providing *Lbx1*^Flpo^ and *Tau*^ds-DTR^ lines. Floxed GRP mice were generated at Johns Hopkins University School of Medicine Department of Neuroscience Murine Mutagenesis Core. The project has been supported by the NIH grants 1R01AR056318-06, R21 NS088861-01A1, R01NS094344, R01 DA037261-01A1 and R56 AR064294-01A1 (Z. F. C).

## Author Contributions Statement

Z.F.C. conceived and directed the project and D.M.B. and Z.F.C. designed the experiments; D.M.B., X.T.L., Q.Y.Y., X.Y.L., X.Z., and B.L.L. performed experiments, D.M.B., and Z.F.C wrote the manuscript.

## Competing financial interests

The author(s) declare no competing financial interests.

## Supplemental Information

Supplemental information includes extended experimental procedures, 8 supplemental figures, 2 supplemental tables and 2 supplemental videos.

## Supplemental Information

### Supplemental Figures S1-S8

**Supplemental Figure 1. Related to Fig 1.**
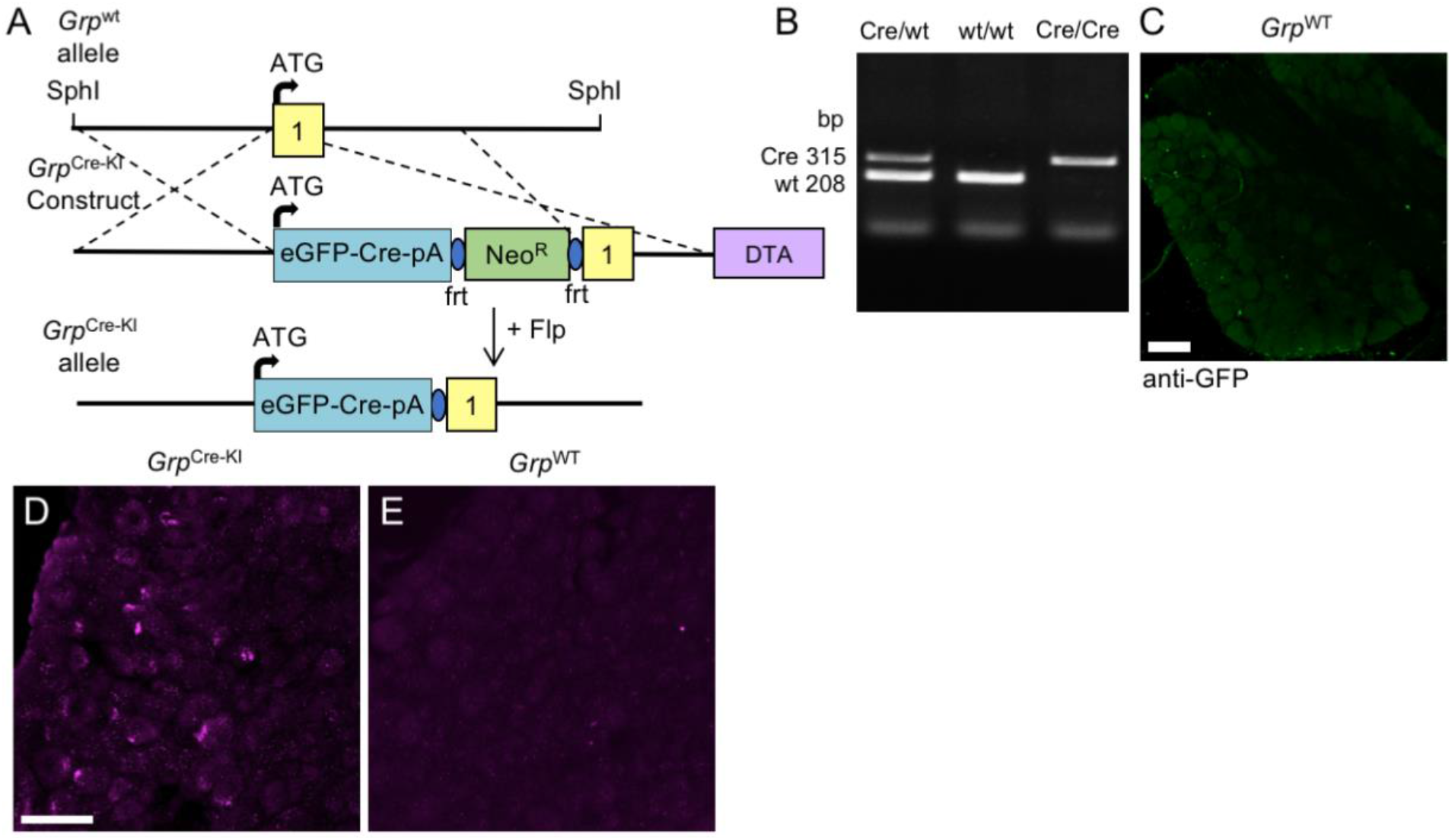
(**A**) Schematic of targeting strategy to knock in eGFP-Cre Neo cassette in *Grp* allele to generate *Grp*^Cre-KI^ mice. (**B**) Gel electrophoresis of genotyping PCR from *Grp* Cre/wt, wt/wt and Cre/Cre samples. (**C**) IHC with eGFP antibody in *Grp*^WT^ DRG. (**D, E**) RNAScope ISH of *Cre* expression in *Grp*^Cre-KI^ (**D**) and *Grp*^WT^ DRG (**E**). Scale bar in **C** and **D**, 50 μm.

**Supplemental Figure 2. Related to Fig 1.**
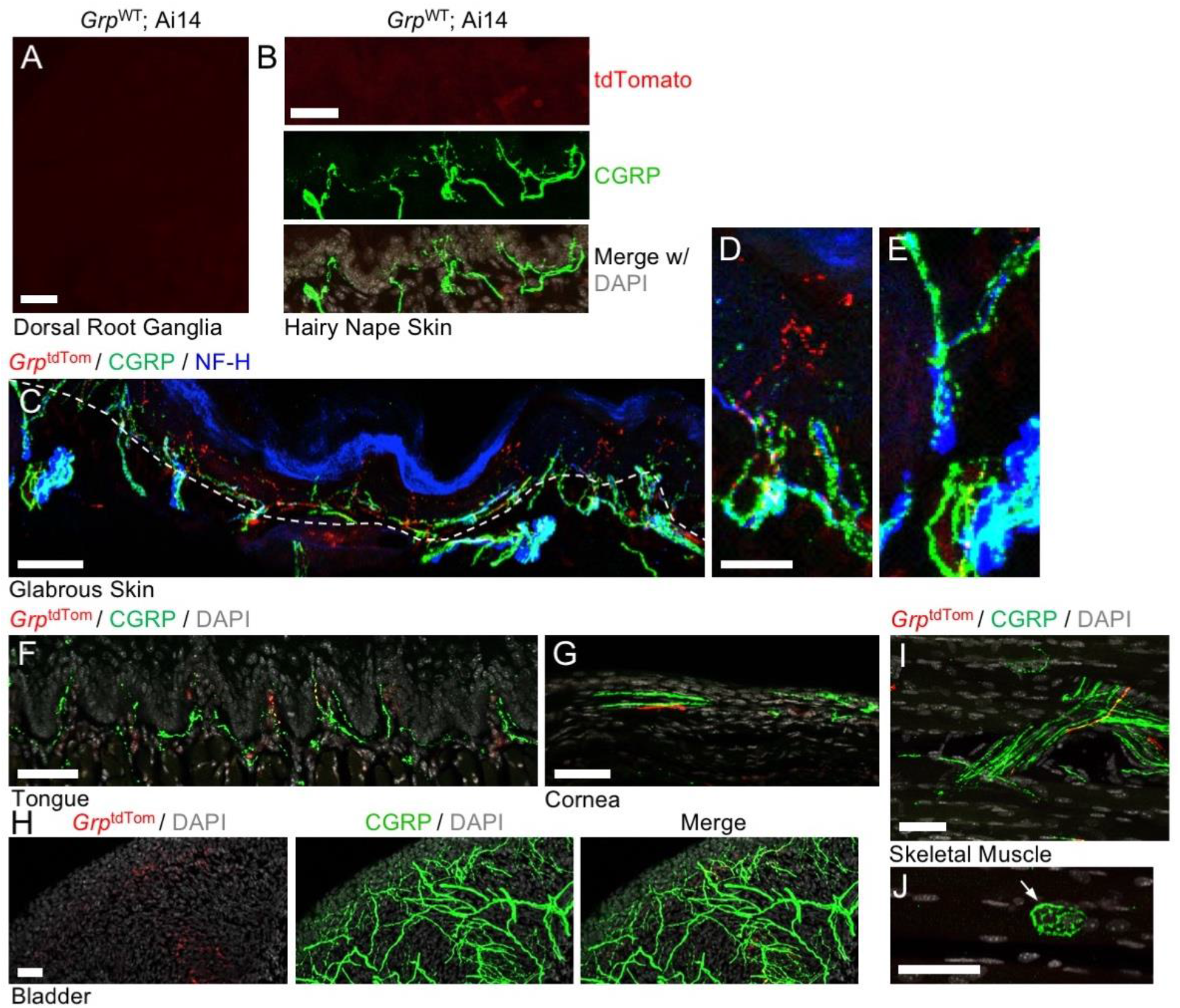
(**A**) Epifluorescent image from *Grp*^WT^; Ai14 DRG. (**B**) IHC images of tdTom, CGRP and DAPI in *Grp*^WT^; Ai14 nape skin. Scale bar in **A** and **B**, 50 μm. **C-E**) IHC images of tdTom, CGRP and NF-H in *Grp*^WT^; Ai14 glabrous skin. Image in **D** shows *Grp*^tdTom^ fiber with no NF-H (blue) expression and **E** shows NF-H fibers that do not express tdTomato. Scale bar in **C** 50 μm and in **D**, 20 μm. **F-J**) IHC images of tdTom, CGRP and DAPI in *Grp*^tdTom^ tongue (**F**), cornea (**G**), bladder (**H**) and skeletal muscle (**I, J**). Arrow indicates neuromuscular junction in l. Scale bar in **F – J**, 50 μm.

**Supplemental Figure 3. Related to Fig 2.**
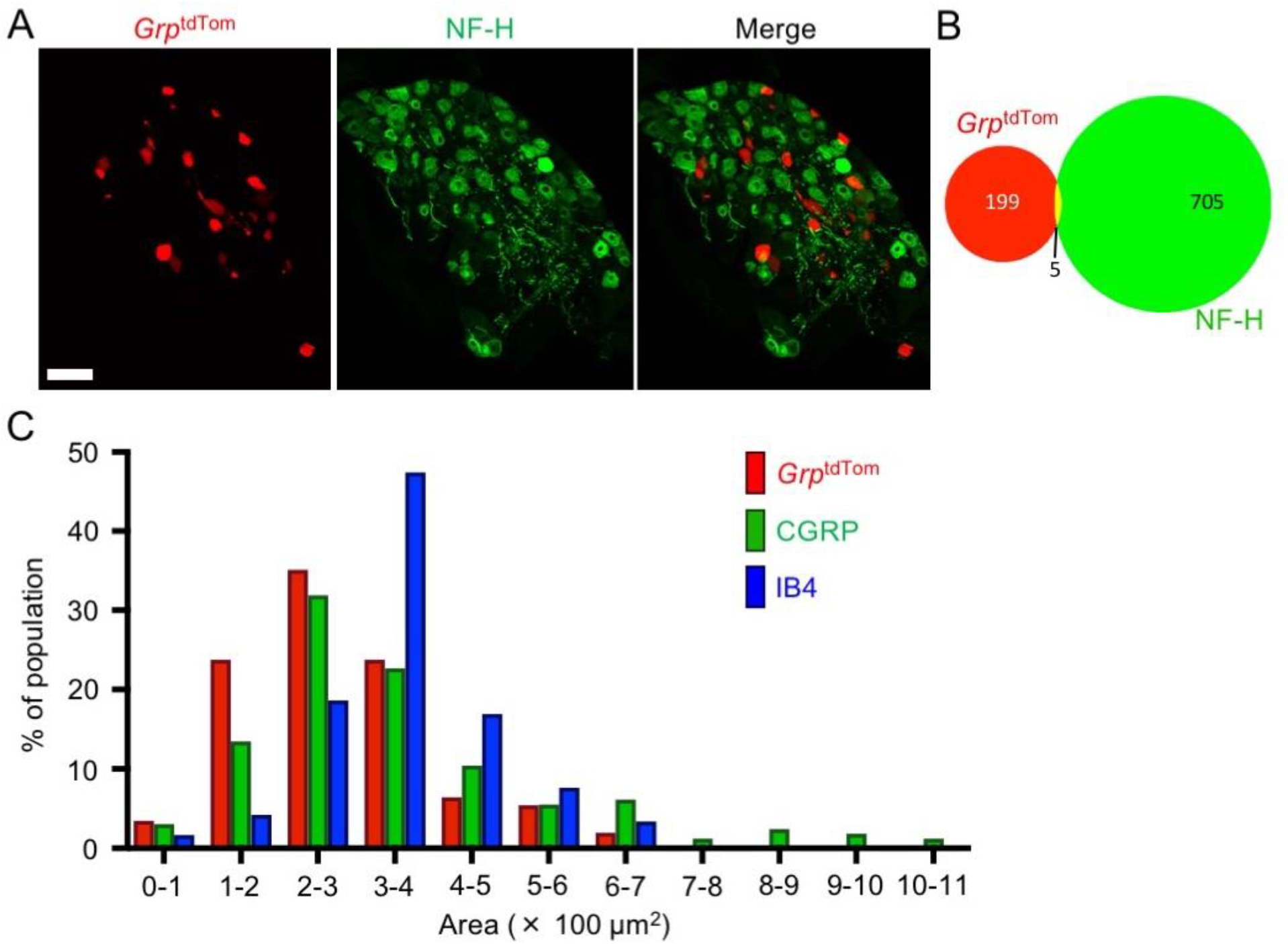
(**A**) IHC of tdTomato and NF-H in *Grp*^tdTom^ DRG. Scale bar in A, 50 μm. (**B**) Diagram of DRG neurons with tdTomato and NF-H expression. (**C**) Frequency Distribution (%) of Perikarya Area (μm^2^) for *Grp*^tdTom^, CGRP or IB4 neuron populations in DRG. Neurons from 6 DRG sections were measured.

**Supplemental Figure 4. Related to Fig 3.**
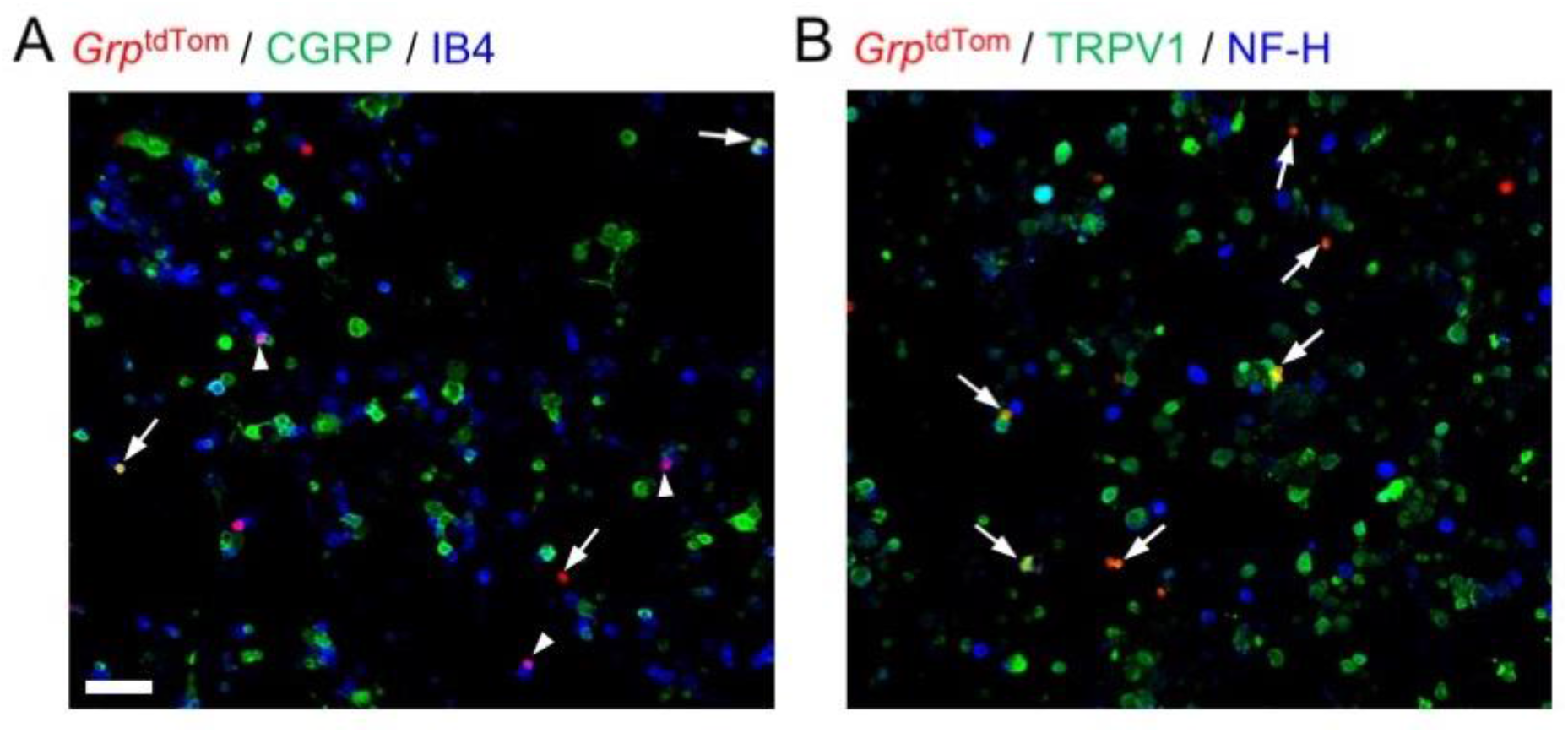
(**A**) IHC image of tdTomato, CGRP and IB4 from *Grp*^tdTom^ DRG cultures. Arrows indicate *Grp*^tdTom^ neurons co-expressing CGRP and arrowheads indicate IB4-binding. (**B**) IHC image of tdTomato, TRPV1 and NF-H from *Grp*^tdTom^ DRG cultures. Arrows indicate *Grp*^tdTom^ neurons co-expressing TRPV1. Scale bar in **A**, 100 μm.

**Supplemental Figure 5. Related to Fig 4.**
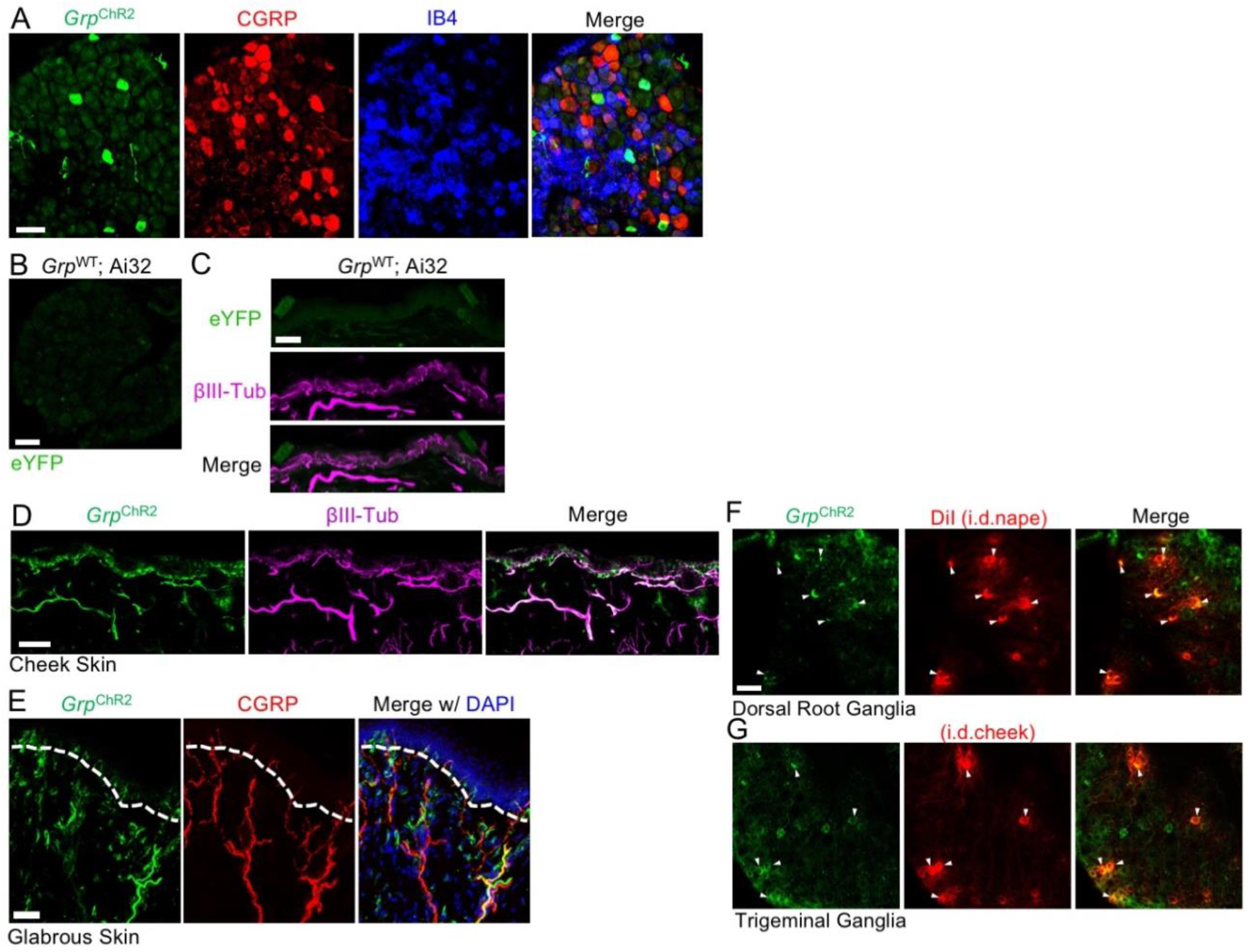
(**A**) IHC images of eYFP, CGRP and IB4 in *Grp*^ChR2^ DRG. Scale bar in **A**, 50 μm. (**B**) eYFP image from *Grp*^WT^; Ai32 DRG. Scale bar in **B**, 50 μm. (**C**) IHC image of eYFP, βIII-Tubulin and DAPI in *Grp*^WT^; Ai32 nape skin. Scale bar in **C**, 100 μm. (**D**) IHC image of eYFP and βIII-Tubulin in *Grp*^ChR2^ cheek skin. Scale bar in **D**, 100 μm. (**E**) IHC images of eYFP, CGRP and DAPI merge in *Grp*^ChR2^ glabrous skin. Scale bar in **E**, 100 μm. (**F**) IHC image of eYFP and DiI in *Grp*^ChR2^ DRG or TG 10 days after i.d. nape or cheek injection of DiI tracer. Scale bar in **F**, 50 μm.

**Supplemental Figure 6. Related to Fig 4.**
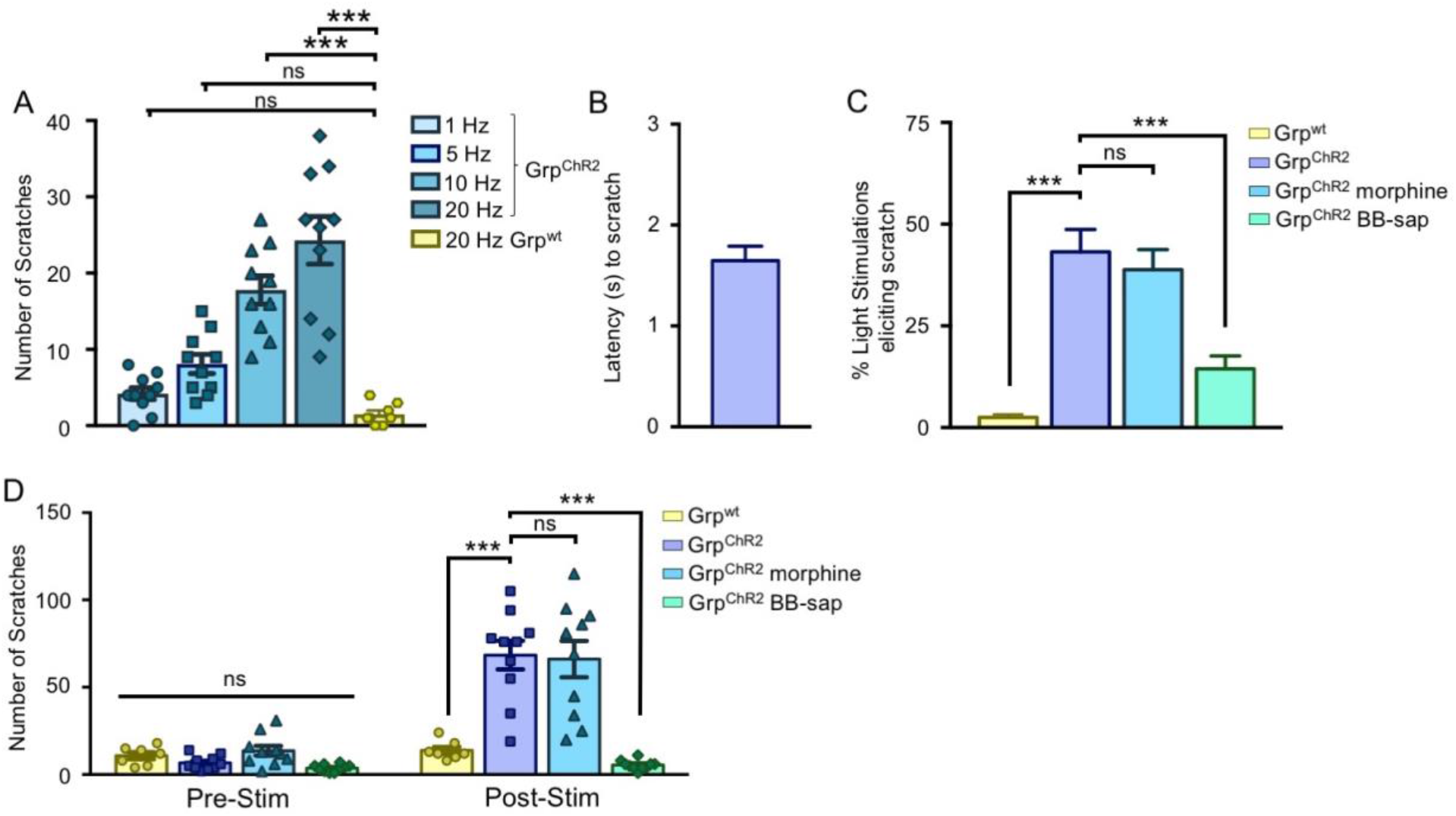
(**A**) Number of scratches in 5 min induced by 3s – 1, 5, 10 or 20 Hz light stimulation of nape skin in *Grp*^ChR2^ and *Grp*^WT^ mice (**B**) Latency to scratch during 3s – 20 Hz light stimulation of skin in *Grp*^ChR2^ mice. (**C**) Percentage of 20 Hz light stimulations eliciting scratching behavior in *Grp*^WT^, *Grp*^ChR2^, *Grp*^ChR2^ morphine-treated and *Grp*^ChR2^ BB-sap-treated mice. (**D**) Number of spontaneous scratches in 30 min before and after the 5-min light stimulation experiment in *Grp*^WT^, *Grp*^ChR2^, *Grp*^ChR2^ morphine-treated and *Grp*^ChR2^ BB-sap treated mice. n = 8 – 10 mice, one-way ANOVA with Tukey *post hoc* in **A** and **C**, twoway RM ANOVA with Tukey *post hoc* in C, *** *p* < 0.001, ns – not significant.

**Supplemental Figure 7. Related to Figure 5.**
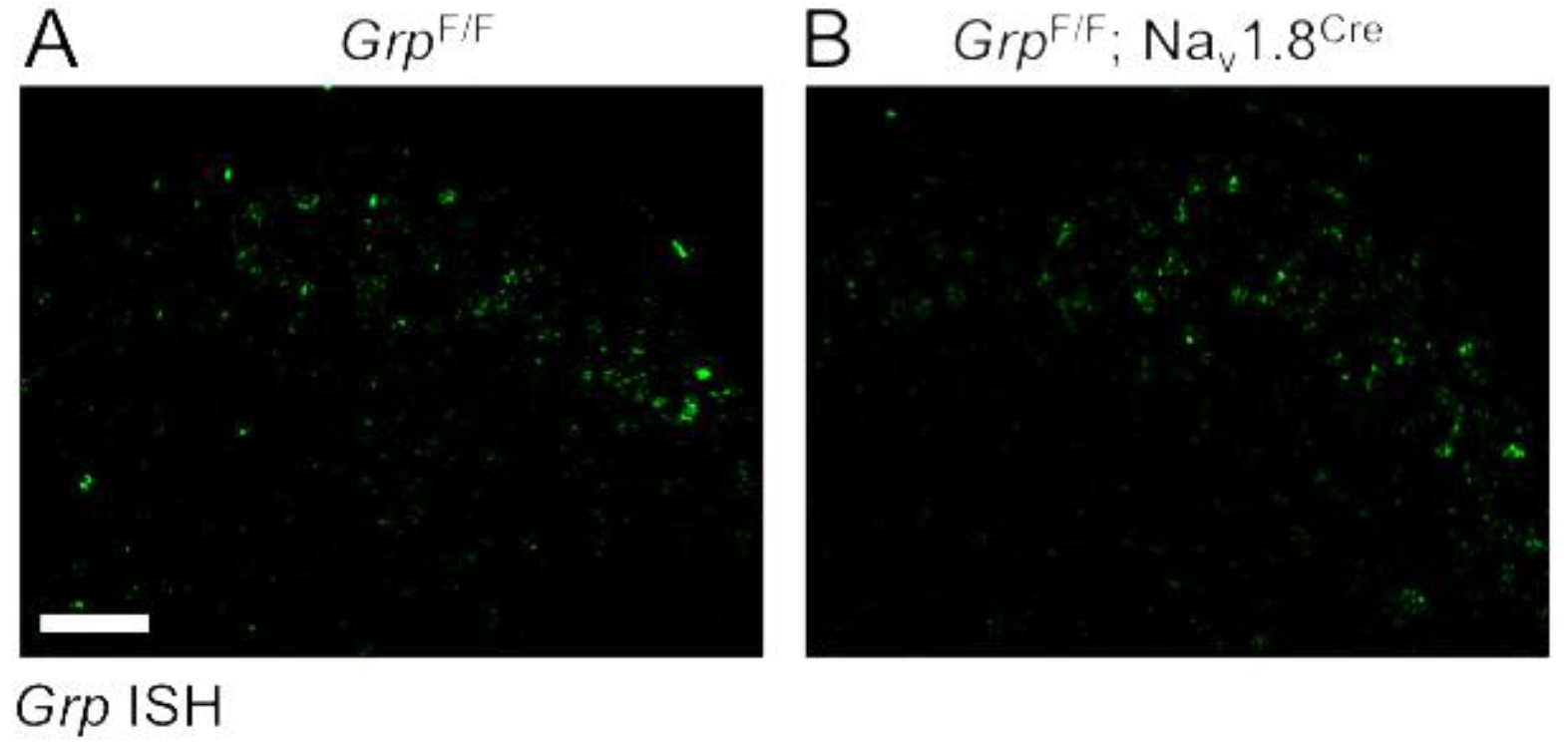
(**A, B**) ISH images of *Grp* expression in spinal cord sections from (**B**) *Grp*^F/F^ and (**C**) *Grp*^F/F^; Na_v_1.8^Cre^ mice. Scale bar in **B**, 100 μm.

**Supplemental Figure 8. Related to Figure 6.**
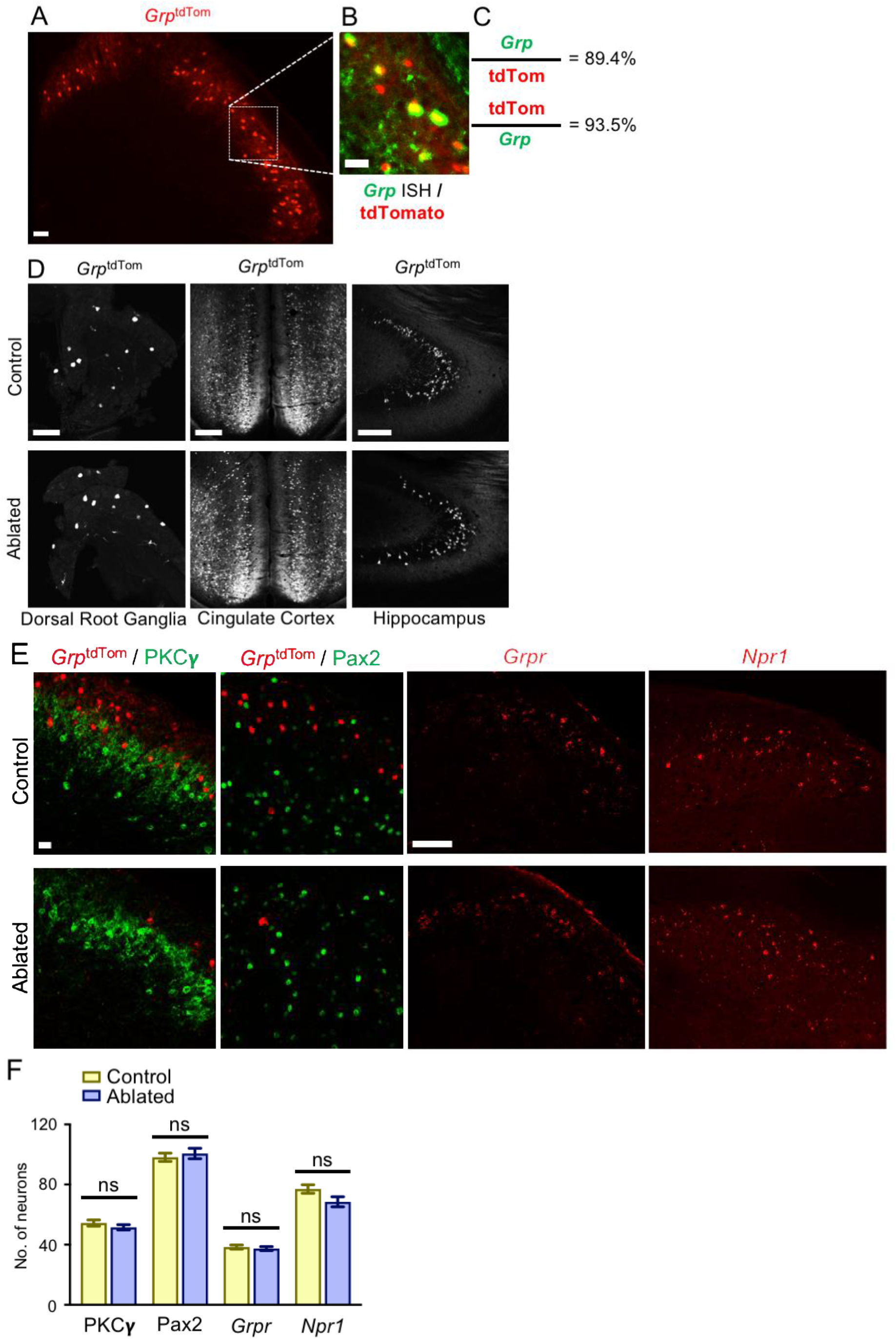
(**A**) Epifluorescent images of tdTomato neurons in cervical spinal cord from *Grp*^tdTom^ mice. (**B**) ISH of *Grp* with tdTomato neurons. (**C**) Percentage of tdTom- or Grp-expressing neurons that express *Grp* or tdTom, respectively. n = 3 mice and 9 sections. (**D**) Epifluorescent images of tdTomato neurons in DRG, cingulate cortex and hippocampus from *Grp*^tdTom^ control and *Grp*^tdTom^ ablated mice. Scale bars in **D**, 50 μm for DRG, 100 μm for cingulate cortex and 100 μm for hippocampus, respectively. (**E**) IHC images of tdTom with PKC*_γ_* or Pax2 and ISH images of *Grpr* or *Npr1* expression in cervical spinal cord sections from *Grp*^tdTom^ control and ablated mice. Scale bars in **E**, 20 μm for tdTom and 100 μm for ISH. (**F**) Mean number of PKC*_γ_*, Pax2, *Grpr* or *Npr1* neurons from control and ablated mice. n = 3 mice and 10 sections, unpaired *t* test in **F**.

### Supplemental Tables S1, S2

**Supplemental Table 1.** Fiber projections in *Grp*^tdTom^ mice

| Tissue | Fibers | Tissue | Fibers |
| --- | --- | --- | --- |
| Skin (Hairy, Glabrous, Mystacial) |  | Retina | +(rare) |
| Unmyelinated: |  | Cornea | +(rare) |
| Epidermal free nerve endings | + | Tongue | +(rare) |
| Circular/penetrating follicle neck endings | + | Bladder | +(rare) |
| Bush/cluster endings | + |  |  |
| Myelinated: |  | Skeletal muscle | -* |
| Circumferential nerve endings | - | Esophagus | - |
| Lanceolate/club endings | - | Heart | - |
| Meissner corpuscles | - | Intestine | - |
| Merkel cells and endings | - | Kidney | - |
| Sebaceous gland endings | - | Liver Lung | - |
| Vibrissa follicle-sinus complex (Vib) | - | Testis | - |
|  |  | Smooth Muscle | - |

\* *Grp<sup>tdTom</sup>* fibers were observed only in nerve bundles going through muscle but did not innervate any muscle fibers.

**Supplemental Table 2.** Single-cell qRT-PCR results from *Grp*^tdTom^ dorsal horn neurons.

| **Gene** | **Percentage (%)** |
| --- | --- |
| *Actb* | 100 |
| *Sst* | 100 |
| *Grp* | 94 |
| *Vglut2* | 85 |
| *Npr1* | 25 |
| *Nmbr* | 12 |
| *Grpr* | 6 |
| *Npy* | 0 |
| *Pdyn* | 0 |
| *Tacr1* | 0 |

n = 4 mice and 10 – 15 neurons

### Supplemental Experimental Procedures

#### Mice

Mice between 7 and 12 weeks old were used for experiments. Ai14 (Stock no. 007908), Ai32 (Stock no. 024109) and C57Bl/6J mice were purchased from Jackson Laboratory, the Na_v_1.8^Cre^ line(Stirling et al., 2005) was kindly provided by the J. Wood lab, and the *Lbx1*^Flpo^ and *Tau*^ds-DJR^ lines (Bourane et al., 2015; Duan et al., 2014) were kindly provided by the Q. Ma and M. Goulding labs. Mice were housed in clear plastic cages with no more than 5 mice per cage in a controlled environment at a constant temperature of ~23°C and humidity of 50 ± 10% with food and water available *ad libitum*. The animal room was on a 12/12 h light / dark cycle with lights on at 7 am. All experiments were performed in accordance with the guidelines of the National Institutes of Health and the International Association for the Study of Pain and were approved by the Animal Studies Committee at Washington University School of Medicine.

#### Generation of *Grp*^Cre-KI^ mice

A ~6kb genomic DNA fragment containing *Grp* exon 1 was PCR amplified from a BAC clone (RP-23, id. 463E10, ThermoFisher Scientific) with a high-fidelty DNA polymerase (CloneAmp HiFi PCR mix, Clontech) and sublconed into pBluescript KS. An eGFP-Cre and frt-flanked PGK-Neomycin cassette for positive selection were integrated into exon 1 ATG start site using an In-Fusion^®^ HD Cloning Kit (Clontech). A diphtheria toxin α (DTA) cassette was inserted downstream of the 3’ homology arm of the construct as a negative selection marker. The construct was linearized with AscI and electorporated into AB1 embryonic stem cells. Following negative selection with G418 (200 ng / μL), positive clones were identified by 5’ and 3’ PCR screening. Positive clones were injected into C57Bl/6J blastocysts to generate chimeric mice. After breeding, germ-line transmission was confirmed by genotype PCR, mice were bred to a FRT-deleter line (Stock. No. 007844, Jackson Laboratory) to remove the Neomycin cassette and were subsequently backcrossed into the C57Bl/6J background.

#### Generation of *Grp* floxed mice

A ~10kb genomic DNA fragment containing *Grp* exon 1 was amplified and loxP sites were integrated into the fragment to flox exon 1. A frt-flanked PGK-Neomycin cassette for positive selection was integrated into intron 1. A diphtheria toxin α (DTA) cassette was inserted downstream of the 3’ homology arm of the construct as a negative selection marker. The construct was linearized with KpnI and electroporated into AB1 embryonic stem cells. Following negative selection with G418 (200 ng / μL), positive clones were identified by 5’ and 3’ PCR screening. Positive clones were injected into C57Bl/6J blastocysts to generate chimeric mice. After breeding, germ-line transmission was confirmed by genotype PCR, mice were bred to a FRT-deleter line (Stock. No. 007844, Jackson Laboratory) to remove the Neomycin cassette and were subsequently bred with Na_v_1.8^Cre^ mice or backcrossed into the C57Bl/6J background.

#### *In situ* hybridization (ISH)

Conventional ISH was performed as previously described (Barry et al., 2016) using a digoxigenin-labeled cRNA (Roche) antisense probe for *Grp* (bases 149-707 of *Grp* mRNA, NCBI accession NM_175012.4). 7 – 12 week-old mice were anesthetized (ketamine, 100 mg / kg and Xylazine, 15 mg / kg) and perfused intracardially with DEPC-treated PBS pH 7.4 followed by 4% paraformaldehyde (PFA) in DEPC-treated PBS. Tissues were dissected, postfixed for 2-4 h, and cryoprotected in 20% sucrose in PBS overnight at 4°C. Cervical, thoracic and lumbar DRGs were sectioned in OCT at 20 μm thickness at 20°C using a cryostat microtome (Leica Instruments) and thaw-mounted onto Super Frost Plus slides (Fisher). Slides were incubated with Proteinase K (50 μg / mL) buffer for 10 min, incubated in prehybridization solution for 3 h at 65°C and then incubated with *Grp* probe (2 μg / mL) in hybridization solution overnight at 65°C. After SSC stringency washes and RNase A (0.1 μg / mL for 30 min) incubation, sections were incubated in 0.01M PBS with 20% sheep serum and 0.1% Tween blocking solution for 3 h and then incubated with anti-digoxigenin antibody conjugated to alkaline phosphatase (0.5 μg / mL, Roche) in blocking solution overnight at 4°C. After washing in PBS with 0.1% Tween, sections were incubated in NBT/BCIP substrate solution at room temperature for 12 – 16 h for colorimetric detection. Reactions were stopped by washing in 0.5% paraformaldehyde in PBS. Bright field images were taken using a Nikon Eclipse Ti-U microscope with a Nikon DS-Fi2 Camera.

RNAScope Double ISH (Advanced Cell Diagnostics, Inc.) was performed as previously described (Wang et al., 2012) using the RNAScope Multiplex Fluorescent Reagent Kit v2 User Manual for Fixed Frozen Tissue. Tissues were dissected, post-fixed for 2-4 h, and cryoprotected in 20% sucrose in PBS overnight at 4°C. Cervical, thoracic and lumbar DRGs or cervical spinal cord regions were sectioned in OCT at 20 μm thickness at 20°C using a cryostat microtome (Leica Instruments) and thaw-mounted onto Super Frost Plus slides (Fisher). Briefly, slides were incubated with hydrogen peroxide for 10 min, washed, mildy boiled in target retrieval reagents for 15 min, washed, dried and hydrophobic barriers were added around the sections. Protease III Plus Reagent was applied for 15 min, washed and sections were incubated with target probes for *Grp* (Mm-Grp-C1, *GenBank:* NM_175012.4, target region bases 22 – 825, Cat No. 317861), *Cre* (Cre-O1-C3, *GenBank:* NC_005856.1, target region bases 2 – 972, Cat No. 474001-C3), *Grpr* (Mm-Grpr-C1, *GenBank:* NM_008177.2, target region bases 463-1596, Cat No. 317871) or *Npr1* (Mm-Npr1-C1, *GenBank:* NM_008727.5, target region bases 941-1882, Cat No. 484531) probes for 2 h. All target probes consisted of 20 ZZ oligonucleotides and were obtained from Advanced Cell Diagnostics. Following probe hybridization, sections underwent a series of probe signal amplification steps followed by incubation of fluorescently labeled probes designed to target the specified channel associated with each probe. Slides were counterstained with DAPI, and coverslips were mounted with FluoromountG (Southern Biotech). Images were taken using a Nikon C2+ confocal microscope system (Nikon Instruments, Inc.) and analysis of images was performed using ImageJ software from NIH Image (version 1.34e) (Schneider et al., 2012). Positive signals were identified as punctate dots and clusters present around the nucleus and/or cytoplasm. For *Grp/Cre* mRNA co-expression, dot clusters of C1 and C3 channels associated within a DRG cell body was counted as double positive, whereas neurons with only C1 or C3 dot clusters were counted as single positive.

#### Immunohistochemistry

IHC was performed as previously described (Barry et al., 2016). Mice were anesthetized (ketamine, 100 mg / kg and Xylazine, 15 mg / kg) and perfused intracardially with PBS pH 7.4 followed by 4% paraformaldehyde (PFA) in PBS. Tissues were dissected, post-fixed for 2-4 h, and cryoprotected in 20% sucrose in PBS overnight at 4°C. For glabrous and mystacial skin, tissues from 3-week-old mice were harvested immediately following anesthesia overdose, fixed in 2% PFA-PBS with 30% (v/v) Picric acid overnight at 4°C and cryoprotected in 20% sucrose in PBS. Tissues were sectioned in OCT at 20 μm thickness for DRG and spinal cord and 30 μm for all other tissues including tongue, cornea, bladder, muscle, intestine, esophagus, heart, testis, liver, lung, kidney and skin. Free-floating frozen sections were blocked in a 0.01M PBS solution containing 2% donkey serum and 0.3% Triton X-100 followed by incubation with primary antibodies overnight at 4°C, washed three times with PBS, secondary antibodies for 2 h at room temperature and washed again three times. Sections were mounted on slides and ~100 μL FluoromountG (Southern Biotech) was placed on the slide with a coverslip. The following primary antibodies were used: rabbit anti-GFP (1:1000, Molecular Probes, A11122), chicken anti-GFP (1:500, Aves Labs, GFP-1020), rabbit anti-CGRP (1:3000, MilliporeSigma, AB15360), guinea-pig anti-TRPV1 (1:800, Neuromics, GP14100), chicken anti-NF-H (1:2000, EnCor Biotechnology, CPCA-NF-H), rabbit anti-βIII-Tubulin (1:2000, Biolegend, 802001), rabbit anti-PKC*_γ_* (1:1000, Santa Cruz Biotechnology, SC-211), rabbit anti-Pax2 (1:300, ThermoFisher Scientific, 71-6000) IB4-binding was performed using an IB4-AlexaFluor 647 conjugate (2 μg / mL, ThermoFisher Scientific, I32450) and DAPI (2 μg / mL, Sigma, D9542) was used as a nuclear stain of cells. The following secondary antibodies were used: Alexa-Fluor 488 conjugated donkey anti-rabbit (1:1000, Jackson Immuno-Research, 711-545-152), Alexa-Fluor 488 conjugated donkey anti-chicken (1:1000, Jackson ImmunoResearch, 703-545-155), Alexa-Fluor 488 conjugated donkey anti-guinea pig (1:1000, Jackson ImmunoResearch, 706-545-148). Cy3-conjugated donkey anti-rabbit (1:1000, Jackson ImmunoResearch, 711-165-152) and Cy5-conjugated donkey anti-chicken 1:1000, Jackson ImmunoResearch, 703-175-155). Fluorescent Images were taken using a Nikon C2+ confocal microscope system (Nikon Instruments, Inc.) and analysis of images for neuron counting was performed using ImageJ software from NIH Image (version 1.34e).

#### Retrograde labeling of DRG/TG neurons from skin

To retrogradely label DRG or TG neurons innervating the hairy nape or cheek skin, the fluorescent tracer 1,1’-Dioctadecyl-3,3,3’,3’-tetramethylindocarbocyanine perchlorate (DiI) (Sigma, 42364) was injected intradermally using a 30 G hypodermic needle. A total of 4 – 20 μL injections of 5% DiI was made within the nape region on the rostral back skin or 2 – 20 μL injections into the cheek skin in *Grp*^ChR2^ mice. After a 10 days to allow for labeling, mice were anesthetized, perfused and DRG and TG were dissected and processed for IHC.

#### DRG culture and Ca2+ imaging

Ca2+ imaging of DRG cultured neurons was performed as previously described (Kim et al., 2016). DRGs from *Grp*^tdTom^ mice were dissected in neurobasal media (Invitrogen) and incubated with papain (30 μL / 2 mL media; Worthington) for 20 min at 37°C. After washing with PBS (pH 7.4), cells were incubated with collagenase (3 mg / 2 mL media; Worthington) for 20 min at 37°C. After washing with PBS, cells were dissociated with a flame-polished glass pipette in neurobasal media. Dissociated cells were collected through a cell strainer (BD Biosciences) to remove tissue debris. Dissociated DRG cells were resuspended with DRG culture media (2% fetal bovine serum, 2% horse serum, 2% B-27 supplement, and 1X glutamine in neurobasal media; Invitrogen) and plated onto poly-ornithine–coated 12-mm-diameter round glass coverslips in a 24-well plate. After 16 – 24 hours, cell culture media were replaced with Ca2+ imaging buffer (140 mM NaCl, 4 mM KCl, 2 mM CaCl2, 1 mM MgCl2, 5 mM glucose, 10 mM Hepes, adjusted to pH 7.4 with NaOH). Fura-2 acetoxymethyl ester (fura-2 AM) (Invitrogen) was diluted to 2 mM stock in DMSO / 20% pluronic acid. The coverslips were mounted on a Warner Instruments recording chamber (RC 26G) perfused with Ca2+ imaging buffer at a rate of ~2 mL / min. An inverted microscope (Nikon Eclipse Ti 10X objective) with Roper CoolSNAP HQ2 digital camera was used for fura-2 Ca2+ imaging after 340 / 380-nm laser excitations (sampling interval, 2 s; exposure time adjusted for each experiment to ~40 ms for 340 nm and to ~30 ms for 380 nm). CQ (1 mM, Sigma), histamine (50 mM, Sigma), capsaicin (10 or 100 nM, Sigma), or KCl (30 mM, Sigma) was applied to *Grp*^tdTom^ DRG cultures to examine Ca2+ responses. Responsive neurons were identified as ROIs and F_340_ / F_380_ ratios were measured using NIS-Elements (version 3.1, Nikon). Prism 7 (version 7.0c, GraphPad) was used to analyze Ca2+ imaging data.

#### Optical Skin Stimulation Behavior

*Grp*^ChR2^ mice and wild-type littermates (*Grp*^WT^) were used for optical skin stimulation experiments. The nape skin was shaved 3 days prior to stimulation in all mice tested. One day prior to the experiments, each mouse was placed in a plastic arena (10 × 11 × 15 cm) for 30 min to acclimate. Mice were videotaped using SONY HDR-CX190 digital video camcorders from a side angle. Mice were recorded for 30 min prior to stimulation for spontaneous scratches. For blue light skin stimulation, a fiber optic cable that was attached to a fiber-coupled 473 nm blue laser (BL473T8-150FC, Shanghai Laser and Optics Co.) with an ADR-800A adjustable power supply. Laser power output from the fiber optic cable was measured using a photometer (Thor Labs) and set to 15 mW from the fiber tip. An Arduino UNO Rev 3 circuit board (Arduino) was programmed and attached to the laser via a BNC input to control the frequency and timing of the stimulation (1, 5, 10 or 20Hz with 10 ms on-pulse and 3 s On – 3 s Off cycle for 5 min). During stimulation, the mouse was traced manually by a fiber optic cable with ferrule tip that was placed 1-2 cm above the nape skin. Following stimulation, the mice were recorded for 30 min. for spontaneous scratches. Morphine (5 mg / kg, i.p.,) was injected 10 min prior to prestimulation spontaneous scratching observations. Bombesin-saporin (200 ng / 5 μL, i.t., Advanced Targeting Systems, Cat No. IT-40) was injected 2 weeks prior to optical stimulation. Videos were played back on a computer for assessments by observers blinded to the animal groups and genotypes. A scratch was defined as a lifting of the hind limb towards the nape or head to scratch and then a replacing of the limb back to the floor, regardless of how many scratching strokes take place between lifting and lowering of the hind limb.

#### Diphtheria toxin receptor-mediated ablation of *Grp* spinal neurons

To ablate DTR-expressing *Grp* spinal neurons for behavioral and histochemical studies, *Grp*^WT^; *Lbx1*^Flpo^; *Tau*^ds-DTR^ control or *Grp*^Cre-KI^; *Lbx1*^Flpo^; *Tau^ds-DTR^* littermates were intraperitoneally injected with diphtheria toxin (DTX, 40μg/kg; Sigma Cat. No. D0564) dissolved in saline at day 1 and then again at day 4. Behavioral, ISH and IHC experiments were performed 2 weeks after DTX injection.

#### Single Cell qRT-PCR

Single-cell qRT-PCR was carried out using Ambion^®^ Single Cell-to-CT^™^ Kit (Life technologies) in accordance with manufactures instructions. Briefly, single tdTomato neurons in laminae I-II of spinal cordslices from *Grp*^tdTom^ mice was identified by red fluorescence under microscope. Negative pressure was applied to the pipette to isolate cytosol of the cell, which was extruded into 10 μl cell lysis/Dnase I solution for RNA extraction and genomic DNA digestion. After reverse transcription (25°C,10 min/42°C, 60 min/85°C, 5 min) target cDNA was pre-amplified for 14 cycles (95°C, 15 sec/60°C, 4min) in the presence of 0.2x pooled TaqMan assays (ThermoFisher Scientific). Diluted pre-amplification products (1:20 in 1x TE buffer) was used for final qPCR reaction (4 μl, 40 cycles of 95°C 5 sec/60°C 30 sec; StepOnePlus, Applied Biosystems) to examine target gene expression. TaqMan assays used are: *Actb*, Mm01205647_g1; *Grp* Mm00612977_m1, Grpr, Mm01157247_m1; *Nmbr*, Mm00435147_m1; *Npr1*, Mm01220076_g1; *Vglut2*, Mm00499876_m1; *Pdyn*, Mm00457573_m1; *Sst*, Mm00436671_m1; *Npy*, Mm01410146_m1; *Tacr1*, Mm00436892_m1. Data were analyzed using StepOne Software (v2.2.2.) with automatic baseline and threshold was set to 0.2.

#### Acute Itch Behavior

Behavioral experiments were performed during the day (0800 – 1500 h) as previously described(Wan et al., 2017). For injections, mice had their nape shaved a day before the experiments. One day prior to the experiments, each mouse was placed in a plastic arena (10 × 11 × 15 cm) for 30 min to acclimate. On the test day, mice were given at least 10 min to get accustomed to recording conditions prior to injections and recordings. CQ (200 μg, Sigma), SLIGRL-NH2 (100 μg, Genscript), BAM8-22 (100 μg, Genscript) or histamine (200 μg, Sigma) dissolved in a volume of 50 μL saline was injected intradermally using a syringe attached to a SS30M3009 – 3/10 cc, 30G × 3/8” needle (Terumo). Immediately after injections, mice were put into rectangular, transparent observation boxes and videotaped from a side angle. The videos were played back on a computer and quantified by an observer who was blinded to the treatment or genotype. A scratch was defined as a lifting of the hind limb towards the nape or head to scratch and then a replacing of the limb back to the floor, regardless of how many scratching strokes take place between lifting and lowering of the hind limb. Only scratches to the injection site were counted for 30 min.

#### Acute Pain Behavior

*Capsaicin cheek behavior*. One day prior to injection, the cheek skin was shaved. On the test day, mice were given at least 30 min to get accustomed to recording conditions prior to injections and recordings. Capsaicin (20 μg) dissolved in a volume of 20 μL saline with 2% ethanol and 2% Tween was injected intradermally to the cheek using a syringe with 30G × 3/8” needle. Immediately after injections, mice were put into rectangular, transparent observation boxes and videotaped from a side angle. The videos were played back on a computer and quantified by an observer who was blinded to the treatment or genotype. A wipe was defined as a singular motion of one forelimb beginning at the caudal extent of the cheek and proceeding in a rostral direction. Only wipes to the injection site were counted for 30 min.

*Thermal sensitivity*. Thermal sensitivity was determined using hotplate (50, 52, or 56 °C), and Hargreaves assay. For the hotplate test, the latency for the mouse to lick its hindpaw or jump was recorded. For the Hargreaves test, thermal sensitivity was measured using a Hargreaves-type apparatus (IITC Inc.). The latency for the mouse to withdraw from the heat source was recorded.

*Mechanical sensitivity*. Mechanical thresholds were assessed using a set of calibrated von Frey filaments (Stoelting). Each filament was applied 5 consecutive times and the smallest filament that evoked reflexive flinches of the paw on 3 of the 5 trials was taken as paw withdrawal threshold.

#### Statistics

Statistical methods are indicated when used. Values are reported as the mean ± standard error of the mean (SEM). Statistical analyses were performed using Prism 7 (v7.0d, GraphPad, San Diego, CA). For comparison of 1, 5, 10 and 20 Hz stimulation-induced scratching in *Grp*^WT^ and *Grp*^ChR2^ mice, a one-way ANOVA with Tukey *post hoc* analysis was performed. For comparison between *Grp*^WT^, *Grp*^ChR2^, *Grp*^ChR2^-morphine, and *Grp*^ChR2^-BB-Sap behavior analysis during light stimulation, a one-way ANOVA with Tukey *post hoc* analysis was performed. For comparison between of *Grp*^WT^, *Grp*^ChR2^, *Grp*^ChR2^-morphine, and *Grp*^ChR2^-BB-Sap behavior analysis prior to and following light stimulation, a two-way repeated measures ANOVA with Tukey *post hoc* analysis was performed. For comparison between *Grp*^F/F^ and *Grp*^F/F^; Na_v_1.8^Cre^ mice acute itch behavior, unpaired *t* test was performed. For comparison between *Grp* control and *Grp* spinal ablated dorsal horn neuron numbers of various IHC and ISH markers, unpaired *t* test was performed. For comparison between *Grp* control and *Grp* spinal ablated mice pre- and post-DTX injection, two-way repeated measures ANOVA with Tukey *post hoc* analysis was performed. Normality and equal variance tests were performed for all statistical analyses. *P* < 0.05 was considered statistically significant.

### Supplemental Movie Captions

**Movie S1.** 20 Hz Light stimulation of nape skin in *Grp*^wt^ control mice.

**Movie S2.** 20 Hz Light stimulation of nape skin in *Grp*^ChR2^ mice.

